# Canonical transcription termination mechanisms explain a minority of operons in cyanobacteria

**DOI:** 10.1101/2025.11.10.687660

**Authors:** Jennifer A. Cascino, K. Julia Dierksheide, Rishi K. Vishwakarma, Yulia Yuzenkova, Paul Babitzke, Katsuhiko S. Murakami, Gene-Wei Li

## Abstract

Cyanobacteria are the most abundant phototrophs and hold potential as a carbon-negative platform for bioengineering applications. However, these efforts have been hampered by limited mechanistic understanding of their gene expression, including transcription termination. Unlike most bacteria, cyanobacteria lack the transcription termination factor Rho, raising the speculation that all transcription ends with intrinsic terminators. Here we show that most transcription units (TUs) in Synechococcus elongatus PCC 7942 are not terminated by known termination pathways. Although many TUs (52%) have unique, well-defined 3′ ends, only a small fraction have features that resemble canonical intrinsic terminators (22%). The noncanonical 3′ ends broadly lacked strong secondary structure, making it unclear how these ends are protected against 3′-5′ exonucleolytic decay. Furthermore, many TUs (46%) have diverse positions of mRNA 3′ ends, suggesting a potentially diffuse termination signal. Finally, we observed a moderate increase in RNA levels downstream of most defined 3′ ends in the absence of the transcription-repair coupling factor Mfd. This finding indicates that Mfd plays a limited, but widespread, role in RNA end formation, potentially through termination of stalled RNAPs. Together, our work reveals unique end architectures of the cyanobacterial transcriptome and suggests that undescribed transcription termination mechanisms are active in the phylum.

**Importance:** Our understanding of bacterial transcription regulation is largely based on model organisms like Escherichia coli and Bacillus subtilis, yet many of these mechanisms appear absent or divergent in cyanobacteria. These differences limit our fundamental understanding of gene regulation and the applied potential of cyanobacteria in sustainable biomanufacturing. To address this gap, we characterized transcription termination in the model cyanobacterium Synechococcus elongatus PCC 7942. We resolve a longstanding question by showing that intrinsic termination alone cannot account for most termination events in this organism. Pervasive transcript ends lacking intrinsic terminator features and the absence of Rho suggest the existence of novel termination mechanism(s) and highlight a largely unexplored regulatory landscape. Simultaneously, our work expands the repertoire of functionally characterized cyanobacterial intrinsic terminators, offering a new toolkit to fine-tune gene expression using terminators of defined strengths. These findings pave the way for more predictable and powerful applications of cyanobacteria in green biotechnology.

## Introduction

Transcription termination is a critical regulatory step in gene expression for generating accurate RNA products and recycling the RNA polymerase (RNAP)^1,2^. The point of termination shapes the 3′ untranslated region and mRNA stability, while the efficiency of termination determines levels of downstream gene expression^3–8^. Furthermore, failure to terminate RNAP can have a range of harmful effects on cells, including decreased production of functional transcripts and increases in antisense transcription or transcription-replication conflicts^1,2,9,10^. Despite this crucial status, the mechanisms for terminating transcription in cyanobacteria—the most abundant group of organisms responsible for carbon fixation on Earth^11^—remain elusive.

In most bacteria, two major termination mechanisms are recognized^1,2^. Named for their respective dependence on the trans-acting termination factor Rho^12^, these pathways are known as Rho-dependent and Rho-independent (or intrinsic) termination. In Rho-dependent termination, the homohexamer Rho binds the nascent transcript at pyrimidine-rich, unstructured regions called Rho-utilization (*rut*) sites^1,2^. The helicase and ATPase activities of Rho dislodge the RNA, collapsing the transcription elongation complex (EC)^1,2^. Intrinsic termination instead relies on two requisite sequence motifs in the RNA—a GC-rich hairpin followed by a uridine-rich region (U-tract)^1,2,13–15^. These features destabilize ECs by pausing RNAP at the U-tract, allowing hairpin folding to trigger EC collapse^1,2,13–15^. Many *in silico* platforms have been designed to predict intrinsic terminators from bacterial genomes^16–33^. High-throughput sequencing of bacterial transcriptomes has also been used for genome-wide identification of intrinsic terminators using RNA 3′ ends^5,8,34–47^.

Contrary to most bacteria, cyanobacteria do not encode a known homolog of Rho^48,49^ and have poorly characterized intrinsic terminators, leaving it broadly unclear how they terminate transcription. Given Rho’s absence, it is commonly proposed that termination occurs only through the intrinsic mechanism in cyanobacteria^49–54^, but this hypothesis has not been tested systematically. Previous studies using RNA sequencing or genomic data showed a varying degree of prevalence for intrinsic terminators, but the definition of intrinsic terminator used in these studies did not always include both a hairpin and a U-tract, which are considered the classic features^41,49,50,55^. To date, only nine putative cyanobacterial intrinsic terminators have been tested in their native context, where they exhibited variable termination efficiencies and often performed worse than *E. coli*-derived or synthetic terminators^48,53,54^. Taken together, the prevalence of intrinsic terminators in the cyanobacterial transcriptome remains unclear.

Understanding transcription termination in cyanobacteria is critical given their significance to global photosynthesis and, increasingly, biomanufacturing. Cyanobacteria are the only bacteria capable of oxygenic photosynthesis, with estimates that they contribute to at least 20-30% of global carbon fixation^56,57^. As such, they have been hailed as potential platforms for industrial-scale, sustainable chemical synthesis that takes advantage of their unique ability to fix atmospheric CO_2_ into other carbon-containing metabolites using water and light^57–64^. However, metabolic engineering of cyanobacteria is hindered by poor characterization of genetic parts like transcription terminators^48,64^.

Accurate mapping of RNA 3′ ends can be used to determine the prevalence of bacterial termination mechanisms. In *E. coli*, for example, high-resolution RNA 3′ end mapping showed that ∼40% of precisely defined transcript termini correspond to intrinsic terminators, while another large portion (∼30%) are formed by Rho-dependent termination^5^. Transcripts insensitive to the Rho inhibitor bicyclomycin had a GC-rich hairpin followed by a U-tract upstream of the 3′ end, consistent with intrinsic terminators. By contrast, bicyclomycin-sensitive transcripts had GC-rich hairpins without a strong U-tract at the 3′ end. These transcripts were interpreted to undergo Rho-dependent termination at diffuse positions downstream, followed by exonucleolytic processing to a stabilizing hairpin at the measured end^5^. Consistent with this model, the sequence downstream of these ends was C-rich and G-poor, resembling a *rut* site signature^1,2,5^. By determining the precise end position of transcripts, high-resolution RNA 3′ end mapping in combination with sequence analysis can thus help reveal the termination mechanism underlying their formation and assess intrinsic terminator prevalence in cyanobacteria.

In this study, we used end-enriched RNA sequencing (Rend-seq)^8^ to map RNA 3′ end architectures at single-nucleotide resolution across hundreds of operons in the freshwater cyanobacterium *Synechococcus elongatus* PCC 7942 (*Syn*). With this proverbial magnifying lens on RNA 3′ ends, we classified the *Syn* transcriptome into two types of transcription units (TUs): those with clearly defined 3′ ends and those with diffuse, multi-position 3′ ends. We found that most TUs in *Syn* (52%) have defined 3′ ends, but diffuse ends are also prevalent (46%). About 43% of TUs with defined ends—only 22% of all TUs—have signatures of intrinsic terminators (a hairpin and U-tract), indicating this termination mechanism is far less prevalent than previously thought. The defined ends lacking intrinsic terminator features do not show strong secondary structure but exhibit a 5′-TG-3′ dinucleotide motif that partially overlaps with the bacterial elemental pause sequence, which encodes a signal for RNAP to isomerize into a transiently paused state^65^. The diffuse ends do not have nearby identifiable intrinsic terminators or other shared sequence features. Finally, we examined the role of other potential termination factors. Interestingly, deleting the bacterial transcription-repair coupling factor Mfd increased apparent transcriptional readthrough at most intrinsic terminators and other defined ends without shifting the position of termination, which may be explained by a role in enhancing termination at or downstream of the observed RNA 3′ ends. Together, our work reveals that intrinsic termination plays a minor role in cyanobacteria, identifies a potential contribution from Mfd, and suggests that undescribed transcription termination mechanisms must operate in this phylum to explain how most operons are punctuated.

## Results

### Mapping RNA 3′ ends reveals defined and diffuse end classes

Rend-seq can simultaneously detect the positions of transcription unit (TU) 5′ and 3′ ends at single-nucleotide (nt) resolution, enabling the precise definition of TU boundaries^8^. To assess the prevalence of intrinsic termination in *Syn*, we measured RNA 3′ end positions in the transcriptome using Rend-seq and characterized the sequence features associated with those ends.

Transcript 3′ ends typically exhibit two main profiles. Some TUs end at a sharply-defined position— concentrated at one or a few bases—producing “defined ends”. Others lack a dominant endpoint and instead display a broad distribution of 3′ ends across many positions, resulting in “diffuse ends”. These profiles reflect the diversity of RNA molecules expressed from a gene and can provide insight into the termination mechanisms that generated them. In *E. coli*, for instance, defined ends can result from either intrinsic termination, where the terminator produces RNA molecules of uniform length, or Rho-dependent termination in which diffusely-terminated RNAs are trimmed by exoribonucleases to a stabilizing hairpin^5^. By contrast, the origins of diffuse TU ends are less well understood. Possible explanations include Rho-dependent (or other factor-dependent) termination without trimming, incomplete exoribonucleolytic decay, or uncharacterized termination mechanisms that do not rely on a single dominant site. Thus, TU 3′ end profiles can inform both the diversity of transcript isoforms and the termination pathway underlying them.

Importantly, Rend-seq can distinguish these TU end profiles. In these data, TUs with defined ends exhibit a sharp peak in 3′-mapped reads (“3′ peak”) followed by an immediate signal drop-off after the peak (**Fig. 1a**; see Materials and Methods for Rend-seq peak definition). By contrast, TUs with diffuse ends show a gradual decrease in signal in the 3′-mapped channel (**Fig. 1a**), or, in some cases, several 3′ peaks (Materials and Methods). To automatically classify the 3′ ends of *Syn* TUs as defined or diffuse, we developed a computational pipeline that scores end profiles based on Rend-seq data (Materials and Methods). As a reference for interpreting the *Syn* data, we applied the same pipeline to previously published Rend-seq datasets from *E. coli* MG1655 (*Eco*) and *B. subtilis* 168 (*Bsu*)^8^—well-characterized, Rho-encoding model organisms.

**Figure 1.**
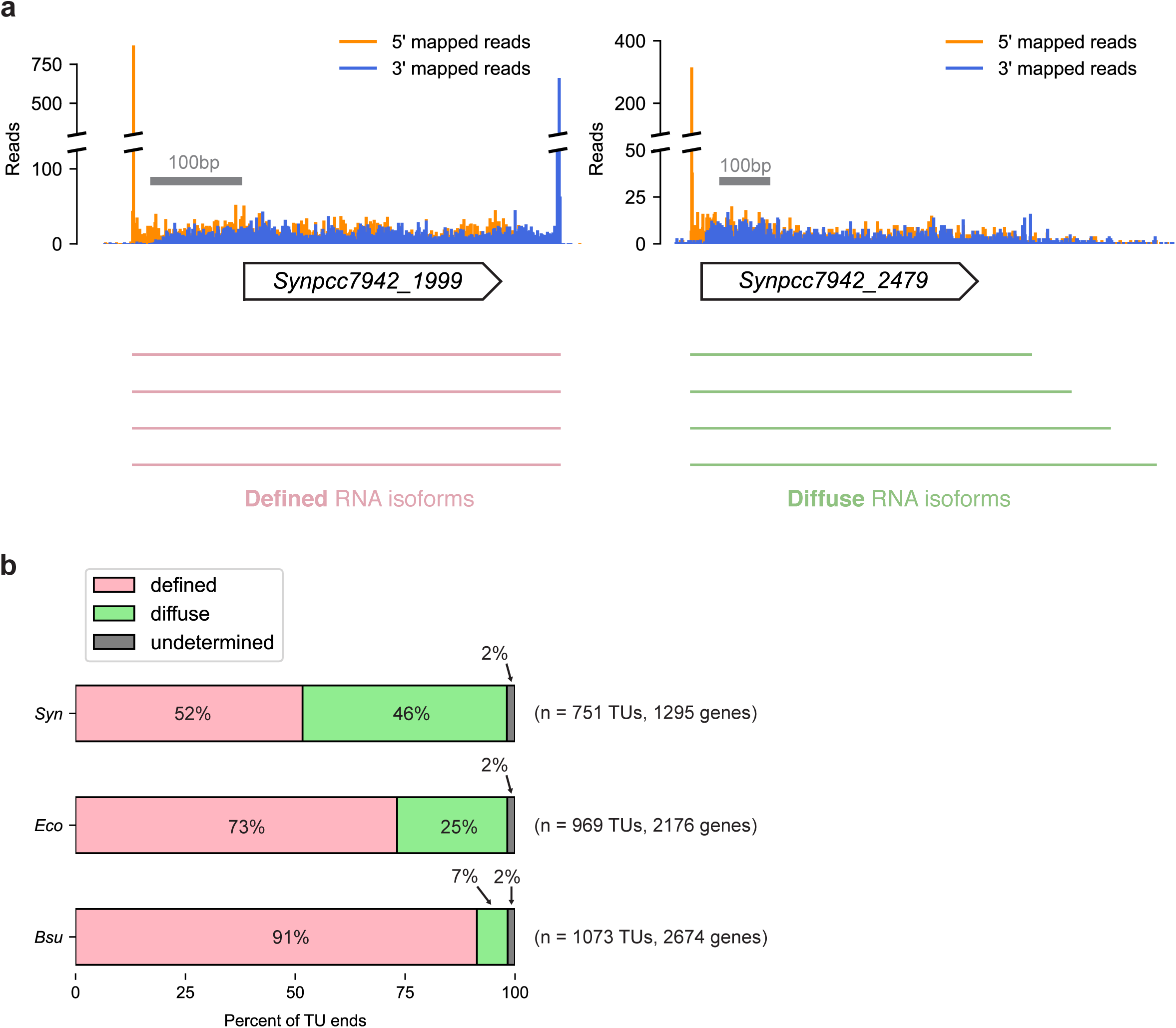
Comparative analysis of TU 3′ end profiles using Rend-seq. (a) Representative Rend-seq traces illustrating defined versus diffuse TU 3′ end profiles. Defined ends exhibit a sharp 3′ peak followed by immediate signal drop-off, corresponding to RNA molecules of uniform length (left), while diffuse ends show a gradual decrease in 3′-mapped reads and lack a dominant end position, corresponding to heterogeneous RNA isoforms (right). Gray bar shows x-axis scale. (b) Genome-wide classification of TU 3′ ends in *Synechococcus elongatus* PCC 7942 (*Syn*), *E. coli* MG1655 (*Eco*), and *B. subtilis* 168 (*Bsu*) using an automated end scoring pipeline (**Materials and Methods**). TUs were classified as having either defined (pink) or diffuse (green) 3′ ends based on parameters in Rend-seq data. End types at a minor subset of TUs could not be determined (gray). Only TUs with sufficient Rend-seq coverage were included. Total number of genes analyzed and corresponding number of TUs (excluding genes internal to TUs) are shown to the right of the bars.

Application of the automatic end scoring pipeline revealed that only a slight majority of *Syn* TUs have defined ends. Of the 751 TUs (1,295 genes) with sufficient sequencing coverage, 52% (n=388) had defined 3′ ends (**Fig. 1b**, Supplementary Table 1). Remarkably, a substantial portion of TUs had diffuse ends lacking a well-defined 3′ end position (46%, n=349). This finding contrasts with the Rho-encoding species, where the vast majority of TUs have defined ends. Indeed, in *Eco* and *Bsu*, defined ends account for 73% (n=709/969) and 91% (n=980/1,073) of TUs, respectively (**Fig. 1b**, Supplementary Table 1). These comparisons are based on similar genome coverage, with the pipeline scoring about half of the annotated genes in *Syn* (n=1,295/2,714), *Eco* (n=2,176/4,651), and *Bsu* (n=2,674/4,546). Thus, in *Syn*, defined 3′ ends account for roughly half of TU ends, whereas diffuse ends are markedly more prevalent than in Rho-encoding bacteria like *Eco* and *Bsu*, suggesting a possible point of divergence in termination dynamics.

### Minority of TU ends are consistent with intrinsic termination

While defined 3′ ends can be produced by both canonical termination pathways, only transcripts punctuated by intrinsic terminators are expected to have a hairpin and a U-tract immediately upstream of the defined end^5^. To assess the prevalence of intrinsic termination in *Syn*, we identified the subset of defined ends with both features.

We found that 58% (n=225) of *Syn* defined ends have ≥4 U in the 8 nt upstream of the 3′ peak, representing 30% of all *Syn* TUs (**Fig. 2a**, **Supplementary Table 2**) (19% of defined ends have 3 U in this region, similar to the probability of 17% by random chance [**Supplementary Fig. 1**]). By comparison, many more putative U-tracts (≥4 U in 8-nt upstream window) were found in *Bsu* (95% of defined ends, representing 86% of all TUs, n=928) (**Fig. 2a**), consistent with reports that intrinsic termination predominates in Firmicutes like *Bsu*, where Rho is dispensable and predicted to play a minor role in terminating coding transcription^22^. We uncovered a similar number of putative U-tracts in *Eco* (49% of defined ends, representing 36% of all TUs, n=346) as in *Syn* (**Fig. 2a**), aligning with evidence that Rho-dependent termination plays a major role in *Eco*, where Rho is essential^22^ and estimated to terminate 20-50% of genes^5^. Together, these data suggest intrinsic terminators are relatively rare in *Syn*, more closely resembling the termination landscape of *Eco*, which relies heavily on Rho, than that of *Bsu*, where intrinsic termination dominates.

**Figure 2.**
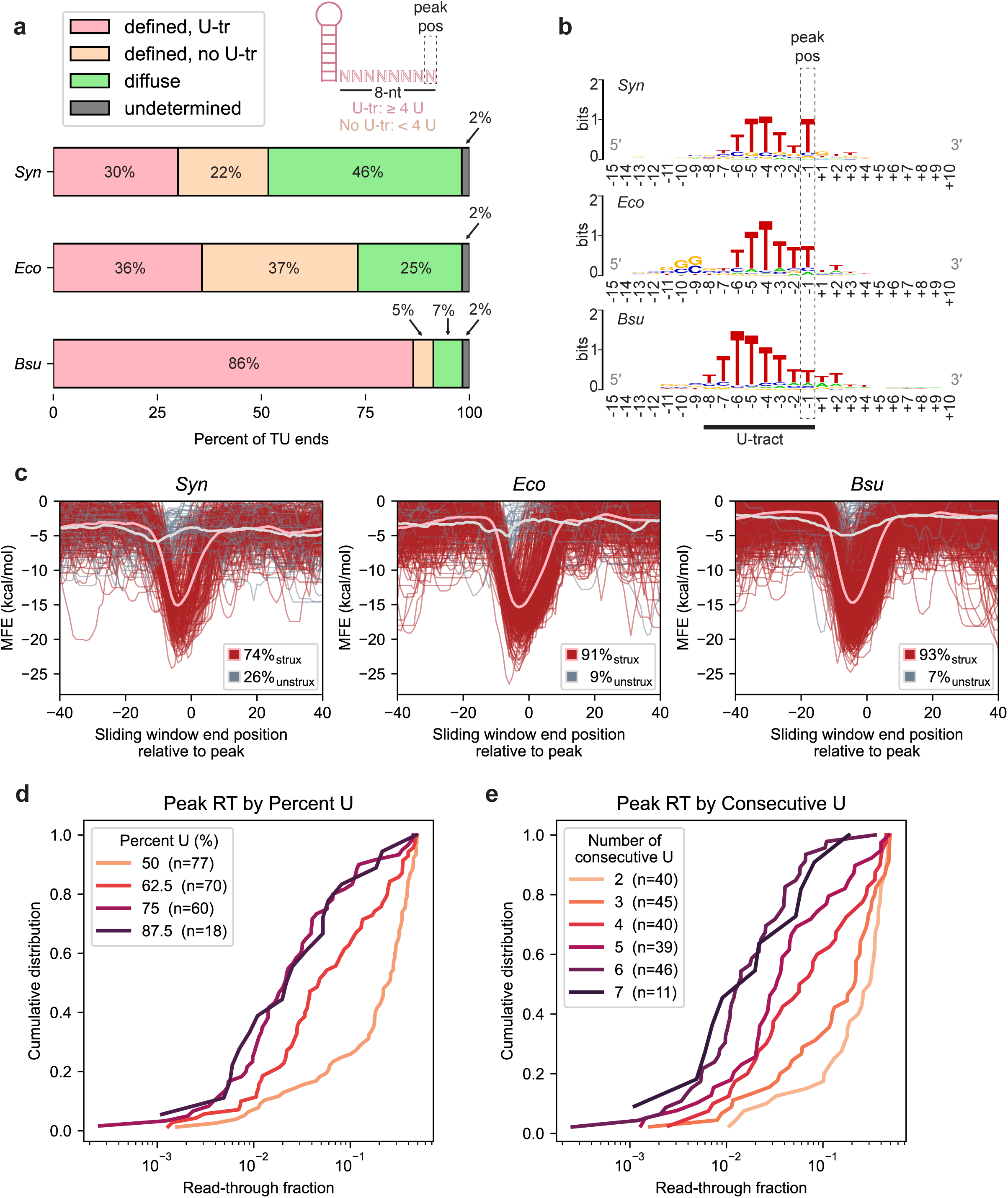
Defined TU 3′ ends with U-tracts are consistent with intrinsic terminators. (a) Proportion of defined TU 3′ ends with (pink) or without (light orange) putative U-tracts in *Synechococcus elongatus* PCC 7942 (*Syn*), *E. coli* MG1655 (*Eco*), and *B. subtilis* 168 (*Bsu*). U-tracts are defined as ≥4 uridines in the 8 nucleotides (nt) directly preceding and including the 3′ peak, which defines the end position. (b) Sequence logos for 25-nt sequence context at defined ends with U-tracts. Peak position and putative U-tract region are shown. (c) Predicted RNA secondary structure at U-tract defined ends. For each end, minimum free energies (MFEs) of 30-nt sliding windows were computed for the 110-nt sequence context of the 3′ peak (**Materials and Methods**). Traces with an MFE < -10 kcal/mol for any of the windows ending from 10 nt upstream of the peak (-10 on x-axis) to the peak itself (0 on x-axis) were colored red to convey the frequency of ends with high RNA secondary structure in the region upstream of the peak where a terminator hairpin is expected to fold. Traces that did not meet this criterion were colored gray to convey the frequency of ends that are not highly structured in the terminator hairpin region. Overlaid pink and light gray traces represent the average MFE trace for ends classified as highly structured (red traces) and not highly structured (gray traces), respectively. Frequency of structured and unstructured ends are listed in inset. (d) Cumulative distribution of readthrough levels at *Syn* U-tract defined ends as a function of U-content (%U) in the putative U-tract region. Number of ends for each category are listed. (e) Cumulative distribution of readthrough levels at *Syn* U-tract defined ends as a function of U-tract length (maximum number of consecutive uridines) in the putative U-tract region. Number of ends for each category are listed. Ends with no consecutive uridines (n=4) have been omitted. For (d-e), readthrough was calculated as described in **Materials and Methods**. Peak pos = 3′ peak (defined end) position; U-tr = U-tract; strux = structured; unstrux = unstructured; RT = transcriptional readthrough.

Analysis of nucleotide information content around U-tract-containing defined ends revealed similar U-rich signatures in *Syn* and *Eco* (**Fig. 2b**). *Bsu* exhibited a more extended U-rich signal in the putative U-tract region (**Fig. 2b**), which suggests that, in addition to their greater prevalence, intrinsic terminators in *Bsu* have more complete U-tracts. In *Bsu*, and to a lesser extent *Eco* and *Syn*, the U-rich signature also extends slightly beyond the defined end. This could reflect how defined end positions were assigned, where closely spaced 3′ peaks (within ±5 nt) are collapsed to the single highest peak (by z-score, **Materials and Methods**). These suppressed minor peaks could represent valid RNA isoforms produced by limited multi-position intrinsic termination. Alternatively, 3′-5′ exoribonucleolytic trimming could shift the measured peak into the U-tract. Notably, although the nucleotides at the 3′ end can be more variable than at other positions of the U-tract in *E. coli*^2^, the U at the -1 (peak) position is strongly enriched in *Syn* (**Fig. 2b**).

We also confirmed that defined ends with U-tracts have strong secondary structure consistent with intrinsic terminator hairpins. Folding 30-nt sliding windows around the 3′ peak at these U-tract-containing ends *in silico* revealed widespread upstream structure in *Syn*, with 74% (n=167/225) having a minimum free energy (MFE) below -10 kcal/mol in the expected intrinsic terminator hairpin region (**Fig. 2c**, **Materials and Methods**). Most *Eco* and *Bsu* U-tract defined ends (91-93%) also had strong secondary structure. Taken together, these analyses suggest that most defined ends with U-tracts in *Syn* are consistent with canonical intrinsic terminators, although defined ends with evidence of both an upstream hairpin and U-tract account for only 22% of all TUs (74% of 30% U-tract-containing defined ends).

To further demonstrate that U-tract defined ends in *Syn* are likely formed by intrinsic termination, we analyzed termination activity at these sites. Since stronger U-tracts are correlated with higher termination efficiency (i.e., reduced transcriptional readthrough) in *Bsu* intrinsic termination^8^, we analyzed the relationship between U-tract composition and readthrough for all 225 *Syn* defined ends with U-tracts (**Materials and Methods**). Ends with higher U-content or more consecutive uridines showed systematically lower readthrough, consistent with stronger termination (**Fig. 2d-e**). Thus, U-tract strength correlates with termination efficiency at *Syn* U-tract defined ends, supporting their function as intrinsic terminators. Finally, we used an *in vitro* transcription system to directly test the termination activity of the putative intrinsic terminators corresponding to two of the U-tract defined ends (**Materials and Methods**). Both sequences tested terminated transcription of *Syn* RNAP, and they were stimulated by NusA, as seen for many intrinsic terminators in *B. subtilis* and *E. coli*^35,66–70^ (**Supplementary Fig. 2**). In sum, these data indicate that only 22% of TUs in *Syn* are intrinsically terminated. Contrary to the prevailing model, canonical intrinsic termination does not appear to be the sole termination mechanism active in cyanobacteria.

### Many defined 3′ ends are inconsistent with intrinsic termination

A fifth of all *Syn* TUs (22%, n=163) have defined ends without U-tracts (**Fig. 2a**, **Supplementary Table 2**). To investigate their origin, we profiled their sequence and structure features and compared them to known termination mechanisms. We also drew a comparison to defined ends lacking U-tracts found for over a third of *Eco* TUs (37%, n=363) and a small fraction of *Bsu* TUs (5%, n=52) (**Fig. 2a**).

First, we determined that defined 3′ ends without apparent U-tracts are not formed by intrinsic terminators immediately downstream, followed by limited 3′-5′ exonucleolytic trimming that removes several nucleotides (i.e., part or all) of the U-tract. Sequence logos of the sequences surrounding these ends revealed no additional uridines downstream of the peak position (**Fig. 3a**). Furthermore, these ends in *Syn* also broadly lack the secondary structure associated with intrinsic termination, with only 46% (n=75/163) showing strong upstream structure (**Fig. 3b**). By contrast, most *Eco* ends exhibited high levels of structure (89%, n=324/363), consistent with known roles of repetitive extragenic palindromic (REP) elements—GC-rich hairpins that stabilize RNA 3′ ends but do not mediate termination^71,72^. We also observed a motif matching the consensus REP element^71,73^ upstream of the peak position in *Eco* (**Fig. 3a**). These structures similarly align with prior observations that Rho-dependent termination followed by exonucleolytic processing to a stabilizing hairpin produces defined TU ends with GC-rich hairpins lacking U-tracts^5^. In *Syn*, however, such stabilizing structures are largely absent. How the majority of these TU ends are formed and stabilized against exonucleolytic decay by PNPase or RNase II in cyanobacteria remains unclear, but local RNA secondary structure does not appear to be the predominant mechanism.

**Figure 3.**
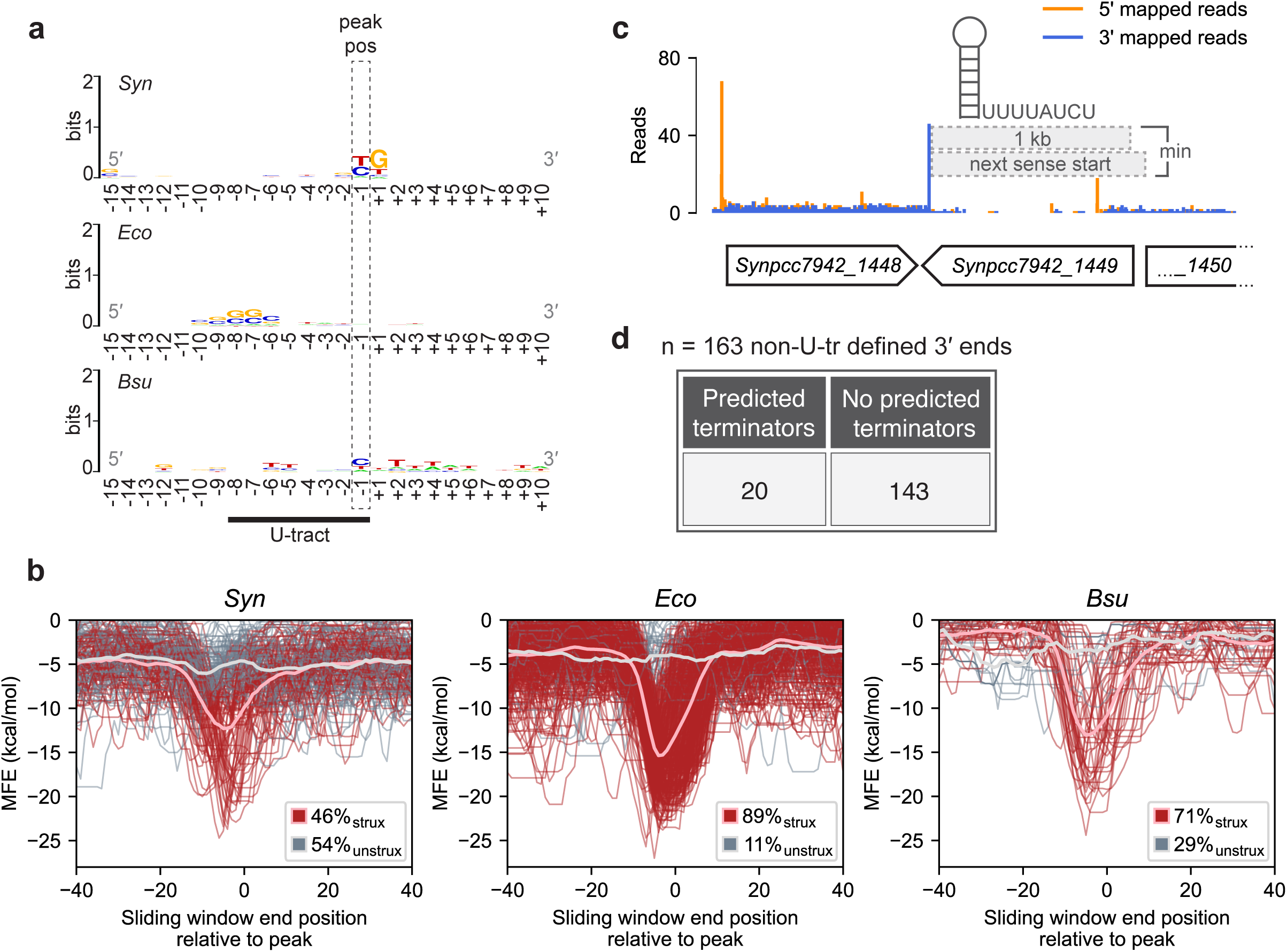
Defined TU 3′ ends without U-tracts are inconsistent with intrinsic termination. (a) Sequence logos for 25-nucleotide (nt) sequence context at defined ends without U-tracts (<4 uridines in 8 nt directly preceding and including the 3′ peak). Peak position and putative U-tract region are shown. (b) Predicted RNA secondary structure at non-U-tract defined ends. For each end, minimum free energies (MFEs) of 30-nt sliding windows were computed for the 110-nt sequence context of the 3′ peak (**Materials and Methods**). Traces with an MFE < -10 kcal/mol for any of the windows ending from 10 nt upstream of the peak (-10 on x-axis) to the peak itself (0 on x-axis) were colored red to convey the frequency of ends with high RNA secondary structure in the region upstream of the peak where a terminator hairpin is expected to fold. Traces that did not meet this criterion were colored gray to convey the frequency of ends that are not highly structured in the terminator hairpin region. Overlaid pink and light gray traces represent the average MFE trace for ends classified as highly structured (red traces) and not highly structured (gray traces), respectively. Frequency of structured and unstructured ends are listed in inset. (c) Analysis of undetected intrinsic termination at non-U-tract defined ends. TransTermHP-predicted intrinsic terminators in the downstream intergenic region of defined ends lacking U-tracts were identified. Only terminators that passed a high confidence filter were included in the analysis (**Materials and Methods**). The intergenic region is defined as the sequence from the stop codon of the gene under consideration to the next start codon on the same strand (limited to 1 kilobase if longer). 1 kilobase gray box is drawn to scale. (d) Frequency of non-U-tract defined ends with a high-confidence intrinsic terminator predicted in the downstream intergenic region. Peak pos = 3′ peak (defined end) position; *Syn* = *Synechococcus elongatus* PCC 7942; *Eco* = *E. coli* MG1655, *Bsu* = *B. subtilis* 168; strux = structured; unstrux = unstructured; *…_1450* = *Synpcc7942_1450*; kb = kilobase; min = minimum; U-tr = U-tract.

Interestingly, we observed a weak sequence motif at *Syn* defined ends lacking U-tracts—a 5′-TG-3′ dinucleotide at the peak and first downstream base (**Fig. 3a**). Overall, the 5′-TG-3′ dinucleotide motif appears at 34% of defined ends without U-tracts (n=56/163). The origin of this motif is unknown, but multiple lines of evidence support its identification as an *in vivo* signature rather than a technical artifact like RNase contamination during library preparation. First, these RNA 3′ ends lack the corresponding downstream 5′ ends that should be formed upon RNase cleavage *in vitro*. Second, 5′-TG-3′ dinucleotide motifs in analyzed *Syn* genes rarely give rise to 3′ peaks—only 0.09% of such instances coincide with a peak (n=80/88,426).

Finally, we assessed if defined ends lacking U-tracts might result from intrinsic termination events farther downstream, followed by extensive RNA processing. Although intrinsic terminator hairpins are considered to protect against exonucleolytic decay, endoribonucleases could separate the terminator from the transcript. Using a program to predict intrinsic terminators (TransTermHP^23^), we predicted putative intrinsic terminators in the *Syn* genome and quantified their presence downstream of genes in each end category (**Fig. 3c-d**, **Materials and Methods**). For genes followed by defined ends lacking U-tracts, only 12% (n=20/163) had a high-confidence intrinsic terminator in the downstream intergenic region, compared to ∼59% of those followed by defined ends with U-tracts (n=132/225). This markedly lower frequency suggests that non-U-tract defined ends are unlikely to arise from undetected intrinsic termination.

The above sequence and structure analyses establish that defined ends without U-tracts in *Syn* are broadly inconsistent with intrinsic termination. These ends lack U-tracts at or immediately downstream of the peak position, and most are not highly structured. Moreover, high-confidence intrinsic terminators are largely absent downstream. We observe a 5′-TG-3′ dinucleotide motif at the peak and +1 position, with evidence supporting its biological origin. However, its role in TU end formation remains unclear, and the broader mechanism(s) that terminate and stabilize these transcripts against decay are not known.

### RNA levels can ramp down long after genes without known terminators

We also uncovered a class of transcript 3′ ends that taper off over many positions indicating RNA isoform heterogeneity. In *Syn*, 46% of TU ends tapered diffusely (n=349), compared to just 25% (n=243) in *Eco* and 7% (n=75) in *Bsu* (**Fig. 2a**). In *Syn*, these ends typically run 100-400 nt until read density drops to ∼30% of the level within the gene body (**Fig. 4a-b**, **Supplementary Fig. 3**, **Materials and Methods**). Only 4% of *Syn* genes with diffuse ends had predicted high-confidence intrinsic terminators downstream (n=14/349) (**Fig. 4c-d**, **Materials and Methods**), indicating that diffuse ends likely are not generated by processing from intrinsic terminators and arise through a distinct mechanism.

**Figure 4.**
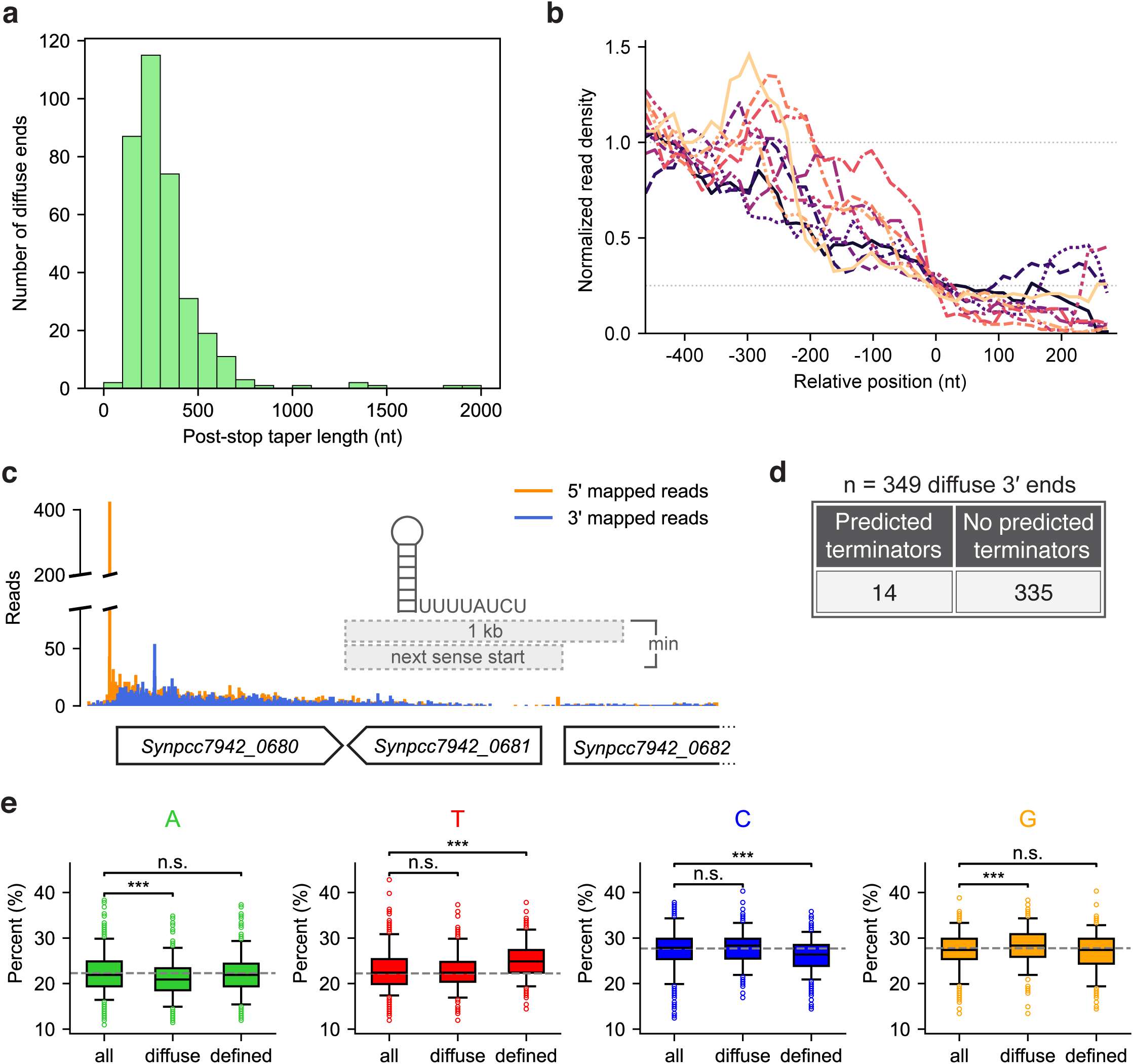
*Syn* diffuse TU 3′ ends taper long after genes without known terminators. (a) Distribution of post-stop tapering lengths for *Synechococcus elongatus* PCC 7942 (*Syn*) diffuse 3′ ends. Diffuse end positions were identified based on a ∼70% drop in read density downstream of the stop codon relative to the upstream coding region (**Materials and Methods**). (b) Mapping diffuse 3′ end tapering in *Syn*. Normalized read densities for a random subset of 10 diffuse ends were mapped for 100-nucleotide (nt) sliding windows around the diffuse end position. Position 0 denotes the window ending at the diffuse end position. (c) Analysis of undetected intrinsic termination at diffuse ends. TransTermHP-predicted intrinsic terminators in the downstream intergenic region of diffuse ends were identified. Only terminators that passed a high confidence filter were included in the analysis (**Materials and Methods**). The intergenic region is defined as the sequence from the stop codon of the gene under consideration to the next start codon on the same strand (limited to 1 kilobase if longer). 1 kilobase gray box is drawn to scale. (d) Frequency of diffuse ends with a high-confidence intrinsic terminator predicted in the downstream intergenic region. (e) Nucleotide content in the 201-nt region upstream of defined and diffuse TU ends (“defined”, “diffuse”) and the 3′ ends of all internal operon genes (i.e., within TUs but not at TU 3′ end), used to control for general gene-end effects (“all”). Box plots show A/T/C/G content distributions; whiskers represent the 5^th^ and 95^th^ percentiles. Gray dotted lines indicate background genomic nucleotide frequencies. Statistical significance was assessed using unpaired two-sided Welch’s t-tests (n.s. = not significant; * = p < 0.05; ** = p < 0.01; *** = p < 0.001). Kb = kilobase; min = minimum.

This tapering profile could result from factor-based termination mechanisms like Rho, which acts over a range of positions^1^. Strikingly, diffuse ends were most abundant in *Syn*, the only species tested that lacks known termination factors to explain this profile. Alternatively, diffuse ends could arise from incomplete exoribonucleolytic decay giving a wide distribution of end positions, or from unknown termination mechanism(s) that do not rely on a single point of termination. Because nucleotide enrichment signatures are known to be associated with shared termination mechanisms like the pyrimidine-rich signature of *rut* sites around Rho-terminated transcript ends^2,5^, we looked at nucleotide preferences at *Syn* diffuse ends (**Materials and Methods**). However, we did not see strong preferences that would help to elucidate their termination mechanism (**Supplementary Fig. 4**), apart from a very modest preference for adenine depletion and guanine enrichment in the 201 nt upstream of diffuse ends (**Fig. 4e**).

### Examination of noncanonical termination factors

Our data demonstrate that most TUs (∼70%) in *Syn* are not terminated by the canonical bacterial mechanisms. However, alternative transcription termination factors have been documented in archaea, eukaryotes, and other bacterial contexts. We therefore investigated whether similar mechanisms could contribute to cyanobacterial termination.

Several termination factors have been reported in eukaryotic organelles of bacterial origin. In chloroplasts, RHON1 is known to mediate some plastid transcription termination in a Rho-like manner^74,75^. However, no RHON1 homologs in *Syn* were found via a protein-protein BLAST. Expanding this search to the entire cyanobacterial phylum returned six low-confidence hits with weak similarity (30-54 aa) to RHON1’s C-terminal region, suggesting they are unlikely to represent functional homologs (**Fig. 5a**, **Materials and Methods**). We also did not detect cyanobacterial homologs of the mitochondrial transcription termination factor MTERF1^76^ (**Fig. 5a**, **Materials and Methods**).

**Figure 5.**
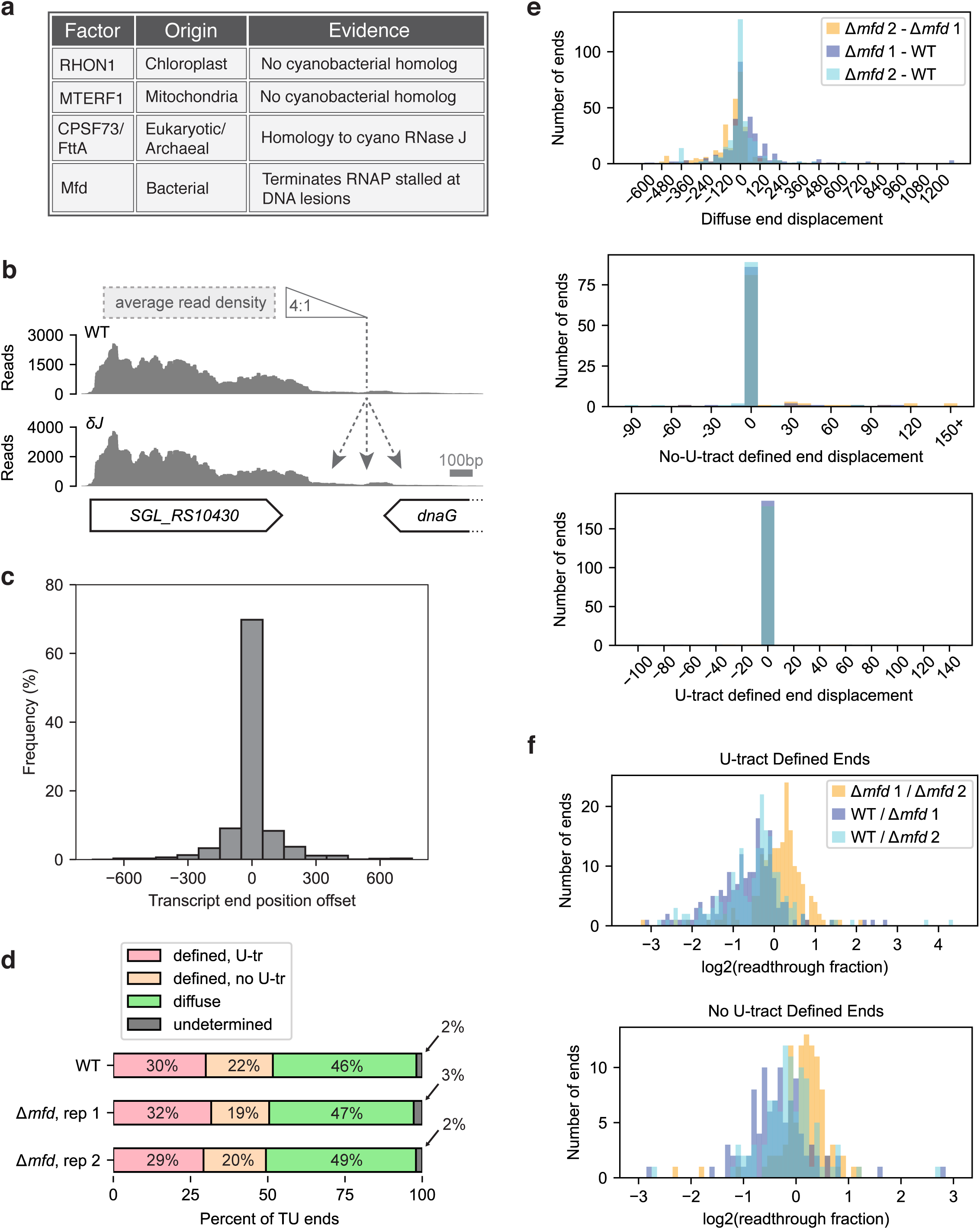
Examination of candidate factor-dependent termination mechanisms in cyanobacteria. (a) Summary of candidate termination factors. Results of BLAST-based homology searches comparing known eukaryotic, archaeal, and organellar factors to the *Synechococcus elongatus* PCC 7942 (*Syn*) genome are listed, as well as rationale for testing bacterial Mfd. (b) RNA-seq data analysis to test for evidence of RNase J termination factor activity using wild-type (WT) and RNase J-depleted *Synechocystis* sp. PCC 6803. Read density in the coding sequence is compared to downstream read densities to identify the window where a drop exceeding 4-fold is achieved, defining the transcript end (**Materials and Methods**). The offset in TU end positions between datasets is calculated as Positiondepletion-PositionWT. Gray bar shows x-axis scale. (c) Distribution of TU end position offsets in RNase J-depleted *Synechocystis* versus WT. (d) Distribution of TU end types identified by automated Rend-seq classification (**Materials and Methods**) for two biological replicates (reps) of Δ*mfd*. End-type distributions were consistent with WT *Syn* for a similar number of genes analyzed (n=1,238 genes, 656 TUs for Δ*mfd* rep 1; n=1,264 genes, 692 TUs for Δ*mfd* rep 2). (e) Distribution of TU end position offsets between WT *Syn* and Δ*mfd* reps for U-tract defined ends, non-U-tract defined ends, and diffuse ends. Offsets were calculated as PositionΔ*mfd*-PositionWT (**Materials and Methods**). (f) Fold-change in transcriptional readthrough at defined ends with or without U-tracts. Readthrough (RT) fractions were calculated as RTWT/RTΔ*mfd* (WT vs. Δ*mfd* reps) or RTΔ*mfd* rep 1/RTΔ*mfd* rep 2 (between reps) (for RT calculation, see **Materials and Methods**). Cyano = cyanobacterial; δJ = RNase J-depleted *Synechocystis*; U-tr = U-tract; WT = wild-type; Δ*mfd* = *Syn* Mfd knockout; Δ*mfd* 1 *=* Δ*mfd* rep 1; Δ*mfd* 2 *=* Δ*mfd* rep 2.

Another candidate is FttA/CPSF73, the conserved termination factor for archaea and eukaryotic Pol II systems^77,78^. Protein-protein BLAST revealed homology of FttA to an ∼100 bp fragment of *Synechococcus elongatus* RNase J (**Fig. 5a**, **Materials and Methods**). More extensive homology to RNase J was identified using DELTA-BLAST, a tool optimized to detect remote protein homologs^79^ (**Materials and Methods**). This 421-bp fragment of *S. elongatus* RNase J shared 16.6% identity to FttA (E value 2x10^-26^) and spanned the full MβL and β-CASPase domains that form the FttA nucleolytic active site^80^. While RNase J was initially identified as a ribonuclease in *Bsu*^81,82^, which encodes paralogs RNase J1 and J2, RNase J1 was recently shown to terminate stalled ECs through a torpedo mechanism that is functionally analogous to FttA/CPSF73-dependent termination^81^.

These findings suggest that RNase J could act as a general cyanobacterial termination factor, mediating an archaeal/eukaryotic-like termination pathway in bacteria. To test this possibility, we reanalyzed RNA-seq data from a *Synechocystis* sp. PCC 6803 RNase J depletion strain^82^. We hypothesized that depletion of a general termination factor would cause widespread transcriptional readthrough leading to downstream shifts in transcript end positions. However, ∼40% of transcripts showed no offset in end position between wild-type (WT) and RNase J-depleted *Synechocystis*, and the overall offset distribution was centered at zero, suggesting any shifts were largely noise-driven (**Fig. 5b-c**, **Materials and Methods**). These findings do not support a role for RNase J as a global cyanobacterial transcription termination factor.

Finally, we assessed whether the transcription-repair coupling factor Mfd contributes to TU end formation in cyanobacteria (**Fig. 5a**). Mfd-mediated transcription termination is the only other well-characterized bacterial termination pathway, but its activity is associated with resolving ECs stalled at DNA lesions genome-wide and not with producing TU ends^1,14,83^. If Mfd were required for termination at TU ends, its loss should cause downstream offsets in transcript end positions relative to WT. To test this possibility, we generated an Mfd knockout strain (Δ*mfd*) and performed Rend-seq on two biological replicates (reps) (**Materials and Methods**, **Supplementary Figs. 4-5**). End-class distributions were consistent with WT for a similar number of genes analyzed (**Fig. 5d**, **Materials and Methods**, **Supplementary Table 1**), and Mfd deletion caused no substantial offsets in TU end positions across classes (**Fig. 5e**). However, there was an overall increase in transcriptional readthrough at defined ends, with the most pronounced effect at those with U-tracts. Among defined ends with U-tracts in both WT and Δ*mfd* datasets for which readthrough fold-changes could be calculated, nearly all showed increased readthrough upon Mfd deletion (**Fig. 5f**, **Materials and Methods**). 25% of these ends (average of replicates) exhibited ≥2-fold increases in readthrough, and half showed at least a 1.35-fold increase (**Fig. 5f**). The effect was less pronounced at defined ends lacking U-tracts (**Fig. 5f**), consistent with their elevated baseline readthrough in WT (**Supplementary Fig. 7**). Thus, Mfd is not required for termination like canonical factors such as Rho, as its loss does not cause complete readthrough or major shifts in end position. Rather, these results could suggest a previously unrecognized role for Mfd in TU end formation. Mfd may act at the observed RNA 3′ ends, potentially stimulating intrinsic termination efficiency, or in the downstream region, where Mfd-dependent termination events are then processed back to the observed endpoints. We did not observe substantial changes in gene expression upon Mfd deletion, so it is unlikely that indirect effects contribute to the changes in readthrough.

## Discussion

Through precision mapping of TU termini, our work reveals that intrinsic termination plays a minor role in terminating transcription in the freshwater cyanobacterium *Syn*. Defined ends in *Syn* that lack intrinsic terminators are not highly structured, leaving their mechanism of formation and stabilization unclear. This contrasts with the predominance of intrinsic terminators in *Bsu* and the prevalence of RNA secondary structures at defined ends in *Eco*. In *Syn*, diffuse 3′ ends—broad, tapering regions that ramp down long after genes without known terminators—were detected at nearly half of all TUs. There were no strong nucleotide biases across these regions, and the functional relevance, if any, of weak preferences remains unknown. Finally, Mfd loss increased apparent transcriptional readthrough at intrinsic sites and, to a lesser extent, other defined ends without shifting the position of termination, suggesting a role in stimulating transcription termination at or downstream of defined ends. The prevalence of TU ends that are not formed by intrinsic termination suggests that additional, unknown termination mechanisms may operate in cyanobacteria.

Mfd is typically associated with an independent termination pathway for removing ECs stalled at sites of DNA damage^1,14,83^. The connection to DNA damage is notable in the context of cyanobacteria, whose photosynthetic lifestyles involve frequent exposure to UV radiation that produces high levels of photodamage to their genomes, including thymine dimers^84,85^. A greater reliance on Mfd’s termination activity in response to elevated DNA lesions could result in the widespread increase in RNA levels we observed downstream of defined 3′ ends upon Mfd deletion (**Fig. 5f**), as the transcript ends generated by Mfd-dependent termination would likely be processed back to stabilizing 3′ ends. Alternatively, it is possible that Mfd acts directly at defined 3′ ends, which may be hotspots of UV-induced thymine dimer lesions due to the enrichment of coding-strand Ts in the U-tract region, consistent with our observation of increased thymine content at defined TU ends (**Fig. 4e**). However, given the low frequency of such lesions (5-40 per megabase of DNA^84^) and the consistency in termination point required to produce a defined end from many uniform-length RNA isoforms, a model whereby Mfd broadly promotes termination and transcript ends are processed back to stabilizing 3′ ends appears more likely. Further characterization of the frequency and position of Mfd-dependent termination will be required to uncover the role of Mfd in shaping transcripts ends in cyanobacteria.

The possibility remains that U-tract-less defined and diffuse TU ends do not reflect transcription termination sites, but rather other processes that can generate transcript 3′ ends. For example, endoribonucleolytic cleavage and/or exoribonucleolytic trimming from downstream termination sites could obscure the true point of termination. In some cases, strong transcriptional pausing alone could appear as a TU end in our data. Additionally, the RNA structures uncovered at 46% of U-tract-less defined ends could be involved in other types of regulation. RNA hairpins are known to stabilize transcripts against decay, guide ribonuclease processing, mediate RNA-RNA interactions, and drive hairpin-stabilized transcriptional pausing^2,65^. Unstructured defined ends and diffuse ends may also be stabilized by small RNAs or RNA-binding proteins. A combination of such processes—e.g., endonucleolytic cleavage followed by selective stabilization—could underlie formation of these non-canonical ends. This possibility represents an important caveat to interpreting the sequence features identified at these ends in our work. Regardless, canonical intrinsic termination remains an unlikely explanation for these end categories given the marked absence of predicted terminators downstream (**Fig. 3d**, **Fig. 4d**).

The 5′-TG-3′ dinucleotide motif common to some defined 3′ ends (**Fig. 3a**) partially overlaps with the consensus bacterial elemental pause sequence—a hairpin-independent pause signal—particularly the Y_-1_G_+1_ motif consisting of a pyrimidine at the nascent RNA 3′ end and guanine at the incoming nucleotide position^65^. Another component of the consensus sequence with a pronounced effect on pausing, G_-10_, is not present in our data^65^. We have several hypotheses for the origin of this motif. First, it could arise from nascent RNA associated with paused RNAPs. Alternatively, these 3′ ends might be generated by pause-induced RNA release that is distinct from previously described termination mechanisms. It could also represent a ribonucleolytic processing signal that may or may not be related to termination.

Several mechanisms could explain termination of TUs with diffuse 3′ ends. First, the RNAP may exhibit reduced processivity in the tapering region, pausing frequently and increasing the likelihood of termination. Additionally, an uncharacterized termination factor could be responsible. One candidate is RNase J, which is homologous to the archaeal termination factor FttA (**Fig. 5a**) that recognizes U-rich RNA^80,86^. Cyanobacteria represent a deep bacterial lineage that branches directly from a common ancestor of Archaea-Eukarya and Bacteria^87^, raising the possibility that a termination factor in this phylum may more closely resemble that of archaea and eukaryotes than the canonical Rho pathway found in many bacteria. We did not observe clear evidence that RNase J acts as a global termination factor in *Synechocystis*; however, the depletion strain was not fully segregated, with PCR confirming persistence of WT *rnj* alleles^82^, suggesting incomplete functional depletion. Clearer RNase J depletion in future studies may uncover stronger impacts on transcription termination.

It is also possible that cyanobacterial RNAP recognizes a non-canonical intrinsic signal, for example, potentially only a U-tract without an accompanying hairpin. It was previously shown that a stretch of uridines is sufficient to pause and induce clamp opening of *Eco* RNAP, which causes nascent RNA release^88^. This reaction was importantly independent of any upstream hairpin structure. Such a signal could account for RNA release at the less structured defined ends with U-tracts (**Fig. 2c**). Another structure-independent pause signal capable of inducing clamp opening and triggering RNA release could also explain defined ends lacking U-tracts, a broadly unstructured end class in our data (**Fig. 3b**), and possibly also diffuse ends.

Practically, our work provides a useful resource for bioengineering^56^. The predictability and efficiency of cyanobacterial metabolic engineering have remained limited, partly due to poorer characterization of genetic parts^48,64^. Our work addresses this gap by vastly expanding the repertoire of functionally characterized intrinsic terminators. We identified 225 defined ends with U-tracts in *Syn* spanning a wide range of predicted efficiencies, including 167 putative terminators that have evidence of both an upstream hairpin and U-tract (**Fig. 2**; see **Supplementary Table 2** for list of these ends and their efficiencies). Although *in vivo* termination efficiency estimates can be influenced by post-transcriptional processes like exonucleolytic trimming^7^, this dataset nonetheless provides a valuable starting point for building tunable, precisely controlled cyanobacterial expression systems.

In summary, the absence of canonical intrinsic terminators in *Syn* reveals a broad, largely uncharted landscape of regulatory biology, potentially shaped by its unique photosynthetic lifestyle, high genomic ploidy, and circadian rhythms. As a lineage that has persisted since life’s earliest days, cyanobacteria represent a largely untapped frontier that is ripe for discoveries that may reshape our understanding of transcription termination and provide new, yet ancient, answers to fundamental questions about the expression codes of life.

## Materials and Methods

### Culturing Conditions

*Syn* WT and Δ*mfd* cells were cultured in BG-11 media optimized for cyanobacteria (Gibco, Catalog No. A1379901), and Δ*mfd* cultures were supplemented with 5 µg/mL kanamycin. Cells were grown at 30°C with 120 rpm of shaking under a 12 hr:12 hr (12:12) light:dark diel cycle with 140-160 µE light intensity (GE 40 Watt, 24” non-dimmable balanced light spectrum LED grow light).

For Rend-seq, 25 mL cultures were inoculated from WT (JAC9) and Δ*mfd* (JAC27) glycerol stocks in 125 mL flasks. When cultures reached ∼mid-exponential phase (OD_730_ = 0.26 [WT] to 0.35 [Δ*mfd*]) 15 days later, they were back-diluted in a total volume of 250 mL in 2.8 L flasks such that 4 doublings were required to reach an OD_730_ = 0.05, the beginning of exponential phase in our growth conditions (target OD_730_ in back-dilution = 0.003125). One flask of WT and two biological replicate flasks of Δ*mfd* (Δ*mfd* reps 1 and 2) were prepared. When cultures had grown up again to exponential phase (4 days later), cells were collected at hour 7 in the 12:12 light:dark diel cycle by pelleting 3 x 35-40 mL of cells at OD_730_ = 0.14-0.22 for 10 min at 4,000 rpm at 4°C (OD_730_ WT = 0.22; Δ*mfd* rep 1 = 0.14; Δ*mfd* rep 2 = 0.19). After pelleting, supernatants were removed, and the cell pellets were flash frozen in liquid nitrogen and stored at -80°C until ready for use in RNA extraction (no longer than 1 month).

### Construction of Mfd Knockout Strain

To knock out endogenous Mfd (*Synpcc7942_1326*), a knockout plasmid was synthesized using TWIST Bioscience’s clonal gene service. A knockout fragment targeting *Synpcc7942_1326* for replacement with a kanamycin resistance (KanR) cassette was inserted into the pTwist Amp High Copy backbone (high copy cloning vector with pMB1 origin of replication and an ampicillin resistance cassette). This knockout fragment contained the promoter and coding sequence of the KanR gene *aphI* encoding aminoglycoside 3′-phosphotransferase derived from the cyanobacterial cloning vector pCV0049 designed by Taton *et al.*^89^, flanked by homology arms targeting the *Synpcc7942_1326* locus. The left and right homology arms comprised the 500 bp sequences immediately flanking the CDS of *Synpcc7942_1326*. The resulting plasmid, pJAC041, was resuspended in 10 mM Tris, pH 8, to a final concentration of 100 ng/µL.

Transformation of pJAC041 into WT *Syn* cells was performed using an established protocol^90^ with slight modifications for lower speed/longer length spins to avoid breaking off transformation pili during cell harvesting as suggested^91^. Briefly, a 25 mL liquid WT *Syn* culture was grown to an OD_730_ of 0.35 under the culturing conditions described in the above section, except constant light was employed, and the culture was used directly after outgrowth from the glycerol stock. A volume of 5 mL of cells was pelleted by centrifugation at 2,000 rpm for 20 min at room temperature. Cells were resuspended in 5 mL of 10 mM NaCl, pelleted by centrifugation as before, then resuspended in 0.3 mL BG-11 and transferred to microfuge tubes. 100 ng (1 µL) of plasmid DNA or 1 µL H_2_O (no DNA control) was added and the tubes were briefly vortexed and inverted several times to mix. Tubes were then wrapped in aluminum foil to block light exposure and incubated in the dark overnight (21 hr) under the culturing conditions described. The next day, the entire cell suspension was plated on a pre-warmed selective BG-11 agar plate (1.5% agar, 5 µg/mL kanamycin) and incubated at 30°C under constant light (∼120-200 µE). Once colonies appeared (6 days later), a patch of transformant cells was restreaked onto a fresh selective plate to ensure full chromosomal segregation. When colonies appeared on that plate (7 days later), a single colony from the second plate was again streaked out onto a new plate. When single colonies appeared on the third plate (7 days later), 5 mL BG-11 liquid cultures supplemented with 5 µg/mL kanamycin were inoculated from single colonies in test tubes. Cultures were grown under the conditions described in the above section, except with constant light. Once cultures were visibly green (6 days later), chromosomal PCR was performed to confirm successful integration of the KanR gene at the *mfd* locus using a previously described protocol^92^. Briefly, 1 mL of culture was pelleted in a microfuge tube at 14,000 g for 4 min at room temperature. The supernatant was removed, and pellets were resuspended in 100 µL TE, transferred to fresh tubes, heated at 100°C for 3 min on a heat block, and centrifuged as before to remove cellular debris. The supernatant was transferred to a new tube, and then 2 µL of cleared supernatant was used in 25 µL PCR reactions with Phusion High-Fidelity DNA polymerase (NEB, Catalog No. M0530L). PCR was performed to detect the left integration junction (primers oJAC259 + oJAC260; 748 bp), right integration junction (oJAC261 + oJAC262; 778 bp), and across the insertion site (oJAC259 + oJAC262; 2,159 bp for the KanR insertion or 4,707 for the WT *Synpcc7942_1326* allele). The cross-site PCR used a 2 min 22 sec extension time to allow amplification of the larger WT product, if present. PCR confirmed successful integration and full chromosomal segregation of the knockout construct (**Supplementary Fig. 5**).

### RNA Extraction

To extract RNA, cell pellets were thawed on ice in 1 mL phenol stop media (6.5:1 BG-11:phenol stop solution; phenol stop solution is composed of 30:18:1:1 100% ethanol:H_2_O:phenol:0.5 M EDTA). One pellet from each Δ*mfd* biological replicate and two WT pellets (technical replicates) were processed, for a total of 4 samples. Thawed pellets were vortexed briefly and centrifuged at 4,000 rpm for 5 min at 4°C, then resuspended in 0.1 mL of TE buffer per 10 mL of original culture at OD_730_ ∼0.2 (350–400 µL TE). NEBExpress® T4 lysozyme (NEB, Catalog No. P8115L) was added at 10 µL per 0.1 mL of resuspended cells (35-45 µL), and samples were incubated at room temperature for 5 min with gentle shaking (300 rpm). Following enzymatic lysis, 2 mL of Buffer RLT (Qiagen) with beta-mercaptoethanol (β-ME) (10 µL β-ME per 1 mL Buffer RLT) were added and samples were vortexed, followed by addition of 2 mL of 70% ethanol. Samples were vortexed, and the resulting ∼4.5 mL lysates were loaded onto RNeasy Midi columns (Qiagen, Catalog No. 75144). RNA was column purified per the manufacturer’s instructions, eluted in 150 µL of nuclease-free H_2_O, and then further purified using a 2x SPRI bead cleanup. Briefly, 2 volumes of SPRI beads were added to each RNA sample, mixed thoroughly by pipetting, and incubated for 5 min at room temperature to allow binding of RNA to the magnetic beads. Samples were then placed on a magnetic rack until the solution cleared (∼5 min), and the supernatant was carefully removed. The beads were washed twice with 1 mL of 80% ethanol on the magnetic rack, incubating for 30 sec each time. Pellets were air-dried on the magnetic rack (∼10 min, until beads appeared slightly cracked), and then RNA was eluted by resuspending beads in a small volume (20-50 µL) of nuclease-free H_2_O, incubating for 5 min at room temperature, and placing on the magnetic rack for 5 min. The supernatant was recovered to retrieve purified RNA. To protect RNA, 1 µL of SuperaseIn RNase Inhibitor (Thermo, Catalog No. AM2696) was added, and samples were then treated with Turbo DNase (Invitrogen, Catalog No. AM2239) per the manufacturer’s protocol to remove contaminating genomic DNA. DNase was heat-inactivated per the manufacturer guidelines, and the reactions were purified by another 2x SPRI bead cleanup as described. RNA quality was confirmed via agarose gel electrophoresis. This protocol effectively lysed cyanobacterial cell walls and yielded high amounts of high-quality RNA (20-30 µg per sample after final cleanup). RNA was stored at - 80°C until Rend-seq library preparation.

### Rend-seq Library Preparation

Rend-seq was performed as previously described^8,93^. rRNA was depleted from 19-20 µg total RNA with MICROBexpress (Invitrogen, Catalog No. AM1905) following the manufacturer’s instructions. Two reactions per sample were performed and then pooled so that the treated RNA amount did not exceed the maximum recommended 10 µg/reaction. Reactions were precipitated in isopropanol. RNA was then fragmented using RNA Fragmentation reagents (Thermo, Catalog No. AM8740) by first incubating RNA in 40 µL of 10 mM Tris, pH 7, at 95°C for 2 min, then adding 4.4 µL 10x Fragmentation Buffer and incubating at 95°C for 25 sec (incubations were performed in a PCR thermocycler). The reaction was quenched with 5 µL Stop Buffer. Samples were precipitated in isopropanol and resuspended in 5 µL 10 mM Tris, pH 7. To size-select RNA, samples were mixed with an equal volume of 2x TBE-urea loading buffer and run on a 15% TBE-urea gel (Thermo, Catalog No. EC6885BOX) for 65 min at 200 V. RNA fragments of 15-45 nt were excised and purified, then precipitated in isopropanol. RNA was dephosphorylated with T4 polynucleotide kinase (NEB, Catalog No. M0201S) in a 20 µL reaction by incubating at 37°C for 1 hr, then the reaction was inactivated at 75°C for 10 min. RNA was precipitated in isopropanol. Linker-1 was then ligated to the 3′ end of 3 picomoles (pmol) of RNA using T4 RNA Ligase 2, truncated K227Q (NEB, Catalog No. M0351L) in a 20 µL reaction containing 25% PEG 8000, T4 RNA Ligase 2 Buffer at 1x, and 100 pmol of Linker-1 by incubating for 2.5 hr at 25°C. Ligated RNA was precipitated in isopropanol and resuspended in 6 µL 10 mM Tris, pH 7, then mixed with an equal volume of 2x TBE-urea loading buffer. The ligated product was then purified by size excision on a 10% TBE-urea gel (Thermo, Catalog No. EC6875BOX) run for 50 min at 200 V and precipitated in isopropanol. cDNA was generated by reverse transcription (RT) with SuperScript III (Invitrogen, Catalog No. 18080093) by first incubating ligated RNA with 25 pmol of RT primer oCJ485 in 12.5 µL reactions at 65°C for 5 min. On ice, the remaining reagents (First Strand Buffer at 1x, 20 U SuperaseIn, 5 mM DTT, 0.5 mM dNTPs, and 1 µL SuperScript III) were then added for a final reaction volume of 20 µL. The reactions were incubated at 50°C for 45 min and then quenched by adding NaOH to a final concentration of 0.1 M and incubating at 95°C for 15 min to hydrolyze template RNA. An equal volume of 2x TBE-urea loading buffer was added, and cDNA was purified by size excision on a 10% TBE-urea gel run for 80 min at 200 V and precipitated in isopropanol. Purified cDNA was resuspended in 15 µL 10 mM Tris, pH 8. Single-stranded cDNA was then circularized with CircLigase (VWR, Catalog No. 76081-606) and additional reagents (2.5 mM MgCl_2_, 50 µM ATP, CircLigase Buffer at 1x) in a final 20 µL reaction volume by incubating at 60°C for 1 hr. Next, 1 µL of CircLigase was spiked in, and the reaction was incubated at 60°C for 1 hr and then inactivated at 80°C for 10 min. The final libraries were amplified by PCR of circularized cDNA and purified by size excision on an 8% TBE gel (Thermo, Catalog No. EC6215BOX) run for 45 min at 180 V. Amplified libraries were precipitated in isopropanol, resuspended in 11 µL 10 mM Tris, pH 8, and stored at -30°C until sequencing.

### Sequencing and FASTQ to WIG File Conversion

Rend-seq libraries were sequenced on an Illumina NextSeq500 platform using 75-nt single-end reads. The FASTQ files returned were converted to WIG files as follows. 3′ linker sequences were stripped, and Bowtie version 1.0.0 (-m 1) was used for sequence alignment of uniquely mapping reads to the reference *Syn* genome CP000100.1. The 5′ and 3′ (end-specific) mapped reads were added separately and strand-specifically at genomic positions. WIG files for the WT technical replicates were combined and processed together in the remaining data processing steps. Shadow-removed WIG files for *Bsu* and *Eco* that were previously generated^8^ were processed alongside the new *Syn* WIG files in the post-shadow removal steps of the data processing pipeline (see “*Data Processing to Generate Shadow-Removed WIG and Peak Sets*” below).

### Peak Calling

Because Rend-seq enriches for the original 5′ and 3′ ends of RNA, TU ends can be mapped at single-nt resolution by identifying sharp peaks in the data. We called Rend-seq peaks slightly differently than previously reported^8^. For each genomic position, two modified z-scores were computed by comparing the read count at that position to a surrounding 50-nt upstream and downstream window, excluding a 5-nt gap on either side to avoid artifactual effects from contiguous peaks (i.e., peak widths > 1 nt). We refer to these as the upstream z-score and downstream z-score. Prior to z-score calculation, read values within the window were winsorized to mitigate the effect of extreme values by replacing outlier values (> 1.5 standard deviations away from the mean) with the mean of non-outliers. The mean (*avg*) and standard deviation (*stdev*) for the winsorized window was calculated, and the modified z-score (*z*) was computed as:

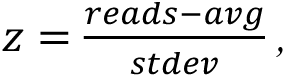

where *reads* refers to the read counts at the genomic position being assessed. Z-scores were only reported for positions with sufficient position-specific read counts and read density in the surrounding window (read threshold ≥ 10 reads, density threshold ≥ 0.25 reads/nt) to avoid false-positives due to counting noise. For these positions, the final z-score was defined as the minimum of the upstream and downstream z-scores. If one of the two could not be computed (e.g., due to insufficient read density in one of the windows), the available z-score was retained. Because the z-score tracks deviation from the local mean signal, true peaks at TU boundaries should be distinguished relative to both flanking regions and have large z-scores in both windows. Taking the minimum ensures that only positions showing a sharp transition in both directions are retained, improving discrimination of true peaks.

Peaks were then called by thresholding on the z-scores. The inverse cumulative distribution (1-CDF) of z-scores was computed, and a threshold was set where the distribution shows a transition between two populations (peaks vs. non-peaks). A z-score threshold of ≥ 12 was used to call peaks on both the original and shadow-removed WIGs for all *Syn* datasets and ≥ 13 on shadow-removed WIGs for *Bsu* and *Eco* (see “*Data Processing to Generate Shadow-Removed WIG and Peak Sets*” below).

### Data Processing to Generate Shadow-Removed WIG and Peak Sets

The data processing pipeline was as follows. First, peaks were called (see “Peak Calling” above) on WIGs in the 5′ and 3′ channel for the three *Syn* datasets (WT, Δ*mfd* rep 1, Δ*mfd* rep 2). Then, the peaks were collapsed. In Rend-seq data, multiple closely spaced peaks may initially be called in the same channel, such that the peak width for a given genomic region—e.g., at a TU boundary—is > 1 nt. At TU ends, clusters of peaks may represent true 3′ ends produced by limited multi-position intrinsic termination or partial 3′-5′ exoribonucleolytic processing that shifts the transcript end by a few nt. To define TU end positions, peaks within ±5 nt were collapsed to the one with the highest z-score. The next step, shadow removal, also requires collapsed peaks as input. Shadow removal was performed as previously reported^8^. Briefly, this process involves removing regions of elevated read density adjacent to peaks (peak “shadows”) that arise as an artifact of Rend-seq end-enrichment. Peaks were then re-called on the shadow-removed WIGs and collapsed again. These final shadow-removed WIG and peak sets were used for TU end calling (see “End Calling Pipeline” below). Note that *Bsu* and *Eco* WIGs were already shadow-removed^8^ and were thus processed starting from the post-shadow removal steps. The following approximate number of reads (in millions, M) were obtained for the *Syn* datasets after processing: 13.5 M (*Syn* WT), 10.2 M (*Syn* Δ*mfd* rep 1), and 10.7 M (*Syn* Δ*mfd* rep 1). Shadow-removed WIG files confirmed the absence of *mfd* transcript in the Δ*mfd* strain, consistent with a successful knockout (**Supplementary Fig. 6**).

### End Calling Pipeline

To automatically identify and classify TU ends in Rend-seq data, we searched the region downstream of each annotated gene (from the annotation file for the complete *Syn* chromosome CP000100.1; *Bsu* chromosome NC_000964.3; and *Eco* chromosome NC_000913.2) for the drop in read density indicative of a transcript endpoint(s). The presence or absence of a single downstream 3′ peak was then used to distinguish between a defined or diffuse transcript end.

To ensure that end classification could be performed confidently, the following criteria were applied. Genes had to be longer than 155 bp, and the winsorized read density (reads above 98^th^ percentile winsorized) across the gene body (excluding 45 bp on either end) had to exceed 0.5 reads/bp. Since 3′ peak detection is limited at low read depth, for genes with a diffuse end classification, we also required that the read depth at the end of the gene (30-105 bp upstream of the stop codon) independently pass a threshold of 0.5 reads/bp. 1,295 genes in WT *Syn*, 2,176 in *Eco*, 2,674 in *Bsu*, 1,238 in Δ*mfd* rep 1, and 1,264 in Δ*mfd* rep 2 were classified. End classification results for all datasets are provided in **Supplementary Table 1**.

Before performing end classification, the set of 3′ peaks identified in each Rend-seq dataset was filtered to generate a set of candidate defined transcription end positions by requiring that read density drop downstream of the peak. For each 3′ peak, the readthrough fraction was determined as previously described^8^. In brief, the readthrough fraction captures the ratio of the read density downstream of the peak to the read density upstream. If a transcript ends at a 3′ peak, the readthrough is expected to be less than 1 due to the drop in read density downstream of the peak. A 3′ peak was therefore included in the set of candidate defined transcription ends if the readthrough fraction was below 0.5.

End classification was then performed as follows. The pipeline alternated between a search for (1) a candidate defined transcription end, and (2) a drop in read density in the region downstream of the stop codon. The search began with a 75-nt window extending 100-175 bp downstream of the stop codon. If the initial downstream window extended into the next TU (marked by a 5′ peak) or past the end of the next gene (marked by a stop codon on the same strand), the regions used for classification were modified as follows. The search for candidate transcription ends was limited to the region between the end of the current gene (10 bp upstream of the stop codon) and the TU boundary (5′ peak or next stop codon position). If no defined end was found, the window for calculating the downstream read density was repositioned to extend 15-90 bp upstream from the TU boundary and trimmed to exclude any overlap with the gene being classified. If the trimmed window was less than 20 bp, the gene was excluded from end classification.

After establishing a downstream window, the region extending from the end of the gene (10 bp upstream of the stop codon) to the end of the downstream window was searched for candidate defined transcription ends. If a candidate defined end was found, a secondary search was performed for any additional 3′ peaks (with any degree of readthrough) between the gene stop codon and candidate defined end. As such additional peaks may represent alternate transcript ends, the end type was recoded to be a “diffuse peak” if additional 3′ peaks were found. In **Figs. 1, 2, 4**, and **5**, the diffuse peaks were counted as diffuse ends. If no additional 3′ peaks were found, an end type of “defined” was assigned to the gene. In some cases, transcription of a gene ended with a 3′ peak, but readthrough could not be defined due to a downstream 5′ peak in close proximity. These genes were classified as having an “undetermined” end type.

If no defined end was found, a diffuse end type was assigned if the read density dropped by at least 3.5-fold between the end of the gene (30-105 bp upstream of the stop codon) and the downstream window. The downstream edge of the window was then recorded as the diffuse end position. If no evidence of a defined or diffuse end was found, the downstream window was stepped downstream by 20 bp, and the two-step search for a defined or diffuse end was repeated with the new window.

This process continued until an end type was assigned, or until the downstream window reached the beginning of the next TU (marked by a 5′ peak) or extended past the stop codon of the following gene on the same strand. At this point, if no evidence of a transcription end was found, the gene was classified as part of a TU (i.e., “TU-internal”).

If at any point the downstream density exceeded the density in the gene by at least 3.5-fold, the end type was assigned to be “undetermined,” as read density is expected to drop or be maintained in the absence of a transcription start site.

### U-tract Analyses

To classify defined ends as having a U-tract, we calculated the percent uridine content in the 8 nt immediately preceding and including the 3′ peak position. U-tracts were scored for peaks with at least 4 uridines (≥ 4 U) in this region. To choose this threshold for U-tract definition, we calculated the number of uridines in randomly sampled 8-nt tracts on both the forward (n=500) and reverse (n=500) strands of the *Syn* genome (n=1000 total tracts) (**Supplementary Fig. 1**). The cumulative distribution of uridine counts in the random 8-nt tracts was plotted. There was a 25% chance of finding ≥ 3 U, 8% chance of ≥ 4 U, and 2% chance of ≥ 5 U. Setting the threshold at ≥ 4 U thus reduced the risk of randomly identifying U-tracts while avoiding excessive stringency.

For the defined ends with U-tracts, we also calculated the maximum number of consecutive uridines in the U-tract. To be counted as having consecutive uridines, at least one ‘UU’ dinucleotide must be found in the 8-nt region. U-tracts with no consecutive uridines must still have 4 uridines to pass the threshold for U-tract definition so will take the form of either ‘UNUNUNUN’ or ‘NUNUNUNU’, where ‘N’ is any non-U ribonucleotide. Such cases (n=4) are omitted from **Fig. 2e**, which shows the remaining 221 U-tract defined ends.

### Sequence Logos

To determine the sequence features surrounding defined ends, we generated sequence logos for the 15 nt upstream of and including the peak position and the downstream 10 nt for defined ends with U-tracts or without U-tracts using WebLogo^94^. Note that WebLogo outputs DNA-based signatures, so T should be interpreted as a U in the RNA.

### Assessing RNA Secondary Structure at Defined Ends

We used Vienna RNAfold (Version 2.4.13)^95^ to calculate the MFE of 30-nt sliding sequence windows starting 70 nt upstream of and including the 3′ peak position to 40 nt downstream of the peak (81 total windows) for all defined ends with or without U-tracts. MFEs for all the windows were plotted against window end positions relative to the peak position, generating a trace that represents the level of RNA structure across the 110-nt sequence context of a given defined end. MFE traces for all defined ends in each U-tract category and species were overlaid to generate a spaghetti plot. Traces with an MFE < -10 kcal/mol for any of the windows ending from 10 nt upstream of the peak (-10 on x-axis) to the peak itself (0 on x-axis) were colored red to convey the frequency of ends with high RNA secondary structure in the region upstream of the peak where intrinsic terminator hairpins are anticipated to fold. Traces that did not meet this criterion in the specified region were colored gray to convey the frequency of ends that are not highly structured in the terminator hairpin region. The overlaid pink and light gray traces on each plot represent the average MFE trace for ends classified as highly structured (red traces) and not highly structured (gray traces), respectively.

Relaxing the MFE threshold to < -9 kcal/mol increased the proportion of structured U-tract defined ends in *Syn* to 77%. Still, 8% of *Syn* ends in this category consistently showed insufficient structure (MFE ≥ -5 kcal/mol) to correspond to a typical intrinsic terminator, compared to just 2% in *Bsu* and 3% in *Eco*.

### Over-Expression and Purification of Cyanobacterial Proteins

Expression and purification of cyRNAP, cy-rpoD1, cyNusG, and cyNusA were performed as follows. All expression plasmids were provided by Dr. Yulia Yuzenkova.

CyRNAP: The cyRNAP core was purified as reported previously^52,96^. In brief, the core enzyme of cyRNAP was overexpressed in *E. coli* BL21(DE3) cells transformed with a pET28a expression vector containing the genes encoding α, β, β′1, β′2, and ω subunits (β and β′2 contain a Strep-tag and His-tag, respectively). The cells were grown in LB media supplemented with kanamycin (50 μg/mL) at 37°C to an OD600 of ∼0.6 then induced with IPTG (1 mM) and grown overnight at 22°C. Cells were harvested by centrifugation and resuspended in lysis buffer (50 mM Tris-HCl, pH 8.0, 250 mM NaCl, 10% glycerol, 1 mM β-mercaptoethanol, and cOmplete^TM^ EDTA-free protease inhibitor cocktail [Roche, Catalog No. 04693132001, used as per manufacturer’s instructions]). The suspension was lysed by sonication (40% amplitude, 5 s on/15 s off cycles, total on-time 5 min), and the lysate was clarified by centrifugation at 18,000 g at 4°C. The supernatant was purified sequentially at 4°C using a HisTrap HP column (5 mL, Cytiva, Catalog No. 17524801) followed by a StrepTrap XT column (1 mL, Cytiva, Catalog No. 29401317). For Ni-affinity purification, the cleared lysate was applied to the HisTrap column pre-equilibrated with Buffer I (50 mM Tris-HCl, pH 8.0, 250 mM NaCl, 2 mM β-mercaptoethanol, and 10% glycerol). The column was washed with Buffer I containing 30 mM imidazole and eluted with Buffer I containing 200 mM imidazole. Eluted fractions were pooled and diluted with 100 mM Tris-HCl, pH 8.0, to adjust the final NaCl concentration to 150 mM for subsequent purification. For Strep-tag affinity purification, the sample was applied to a StrepTrap XT column pre-equilibrated with Buffer W (100 mM Tris-HCl, pH 8.0, 150 mM NaCl, 1 mM EDTA), washed with 3-4 column volumes of Buffer W, and eluted with Buffer E (100 mM Tris-HCl, pH 8.0, 150 mM NaCl, 1 mM EDTA, and 2.5 mM desthiobiotin). The purified cyRNAP was analyzed by SDS-PAGE, concentrated using an Amicon Ultra centrifugal filter unit (100 kDa MWCO, Catalog No. UFC8100), and buffer-exchanged into storage buffer (40 mM Tris-HCl, pH 8.0, 200 mM KCl, 1 mM EDTA, 1 mM DTT, and 5% glycerol). The enzyme was mixed with an equal volume of 100% glycerol, snap frozen in liquid nitrogen, and stored at -80°C until use.

Cy-rpoD1: Recombinant rpoD1 was purified from *E. coli* BL21(DE3) cells carrying the plasmid pET28a-TEV-rpoD1 following a previously described protocol^52^ with minor modifications. Cells were grown in LB medium supplemented with kanamycin (50 µg/mL) at 37°C until reaching an OD_600_ of ∼0.6. Protein expression was induced with 0.4 mM IPTG, and the culture was incubated for 16 hr at 18°C. The harvested cells were resuspended in lysis buffer (50 mM Tris-HCl, pH 7.9, 300 mM NaCl, 5% [v/v] glycerol, 2 mM β-mercaptoethanol, and protease inhibitor cocktail) and lysed by sonication. The clarified lysate was applied to a HisTrap HP column (1 mL, Cytiva, Catalog No. 29051021) pre-equilibrated with Buffer I (50 mM Tris-HCl, pH 7.9, 300 mM NaCl, 2 mM β-mercaptoethanol, and 5% glycerol) containing 10 mM imidazole. The column was washed with Buffer I containing 30 mM imidazole and eluted with Buffer I supplemented with 400 mM imidazole. The eluted fractions were treated with TEV protease during dialysis against buffer (20 mM Tris-HCl, pH 7.8, 300 mM NaCl, 5% [v/v] glycerol, and 2 mM β-mercaptoethanol) to remove the His-tag. The sample was then applied again to a HisTrap HP column (1 mL, Cytiva, Catalog No. 29051021) pre-equilibrated with buffer I, and flow through was collected. The sample was diluted to reduce the NaCl concentration to 100 mM and then applied to a HiTrap Q HP anion exchange column (1 mL, Cytiva, Catalog No. 29051325) pre-equilibrated with 10% Buffer B (Buffer A: 20 mM Tris-HCl, pH 7.9, 0.2 mM EDTA, 1 mM DTT, and 5% glycerol; Buffer B: Buffer A with 1 M NaCl). The bound protein was eluted using a linear salt gradient between Buffer A and Buffer B. Fractions containing rpoD1 were pooled, concentrated using an Amicon Ultra-4 centrifugal filter unit (3 kDa MWCO; Merck Millipore, Catalog No. UFC8003), and buffer exchanged into storage buffer (20 mM Tris-HCl, pH 7.9, 200 mM NaCl, 5% glycerol, 1 mM DTT, and 0.2 mM EDTA). The purified protein was mixed with an equal volume of 100% glycerol, snap frozen, and stored at -80°C until use.

CyNusG and CyNusA: For the preparation of cyanobacterial NusA and NusG, *E. coli* BL21(DE3) cells were transformed with the respective expression plasmids and grown in 500 mL LB medium at 37°C to an OD_600_ of ∼0.5-0.6. Protein expression was induced with 0.5 mM IPTG, and the cultures were incubated for 4 hr at 30°C. Cell pellets were resuspended in 50 mL lysis buffer (40 mM Tris-HCl, pH 7.9, 300 mM NaCl, 2 mM β-mercaptoethanol, and 5% glycerol) supplemented with one tablet of cOmplete^TM^ EDTA-free protease inhibitor cocktail (Roche, Catalog No. 04693132001) and 0.1 mg/mL lysozyme. Cells were lysed by sonication as described above, and the lysate was clarified by centrifugation. The cleared lysate was applied to a HisTrap HP column (1 mL, Cytiva, Catalog No. 29051021) pre-equilibrated with Buffer I (40 mM Tris-HCl, pH 7.9, 300 mM NaCl, 2 mM β-mercaptoethanol, and 5% glycerol). The column was washed with Buffer I containing 30 mM imidazole, and bound proteins were eluted with Buffer I supplemented with 250 mM imidazole. Fractions containing NusA or NusG were pooled and concentrated using Amicon Ultra centrifugal filters. NusA was subjected to an additional purification step prior to gel filtration. Namely, the salt concentration of the pooled NusA fractions was adjusted to 100 mM, and the sample was applied to a HiTrap Q HP anion exchange column (1 mL, Cytiva, Catalog No. 29051325) pre-equilibrated with 10% Buffer B (Buffer A: 20 mM Tris-HCl, pH 7.9, 0.2 mM EDTA, 1 mM DTT, and 5% glycerol; Buffer B: Buffer A containing 1 M NaCl). NusA was eluted using a linear salt gradient between Buffer A and Buffer B. Fractions containing NusA were pooled and concentrated using Amicon filters. Both NusA and NusG were further purified by size-exclusion chromatography on a HiLoad 16/600 Superdex 75 column (Cytiva, Catalog No. 28989333) equilibrated with storage buffer (20 mM Tris-HCl, pH 7.9, 200 mM NaCl, 0.2 mM EDTA, 0.5 mM DTT, and 5% glycerol). Fractions containing pure NusA or NusG were pooled, concentrated, mixed with an equal volume of 100% glycerol, snap frozen in liquid nitrogen, and stored at -80°C until use.

### DNA Templates for In Vitro Transcription Termination Assay

DNA templates for the multi-round transcription assay were synthesized as gene blocks (Integrated DNA Technologies) and PCR amplified using forward and reverse primer sets. Each DNA template contained a promoter region with canonical T7A1 motifs (–10/–35 elements), a transcription start site, and a terminator region comprising either a short (Term1, T-1849) or long (Term2, T-0252) stem-loop structure, followed by an additional short downstream sequence beyond the termination site. Sequences for the two tested terminator templates and amplification primers were: Synpcc_term1 (T-1849), 5′-CGGAATTCCGAAGA TTAATTTAAAATTTATCAAAAAGAGTATTGACTTAAAGTCTAACCTATAGGATACTTACAGCCATATGAGAGTTGCTAGTCGCTTCCCTGTCCAGTCCTCCTGTCTGTCTTAAGCAGCTTAGGCGCGATCGCCTAGGCTGTTTTTTTGACCGTCCATTGCGTGATCGGTGCCAGCGATTTCCAGCCAAGCTTGGG-3′; Synpcc_term2 (T-0252), 5′-CGGAATTCCGAAGATTAATTTAAAATTTATCAAAAAGAGTATTGACTTAAAGTCTAACCTATAGGATACTTACAGCCATATGAGAGTTGAAACCCGCCGATACAGAGGCGTACTATTGCGGCTAGAGTACCGCGGCTCAACCGATGCCAAGGTCTCTAGCTGTTTTTGTGCTTGGCTGCGGGAGCAATCCAGAAGCTGTGCGGCCAAGCTTGGG-3′; Synpcc_F, 5′-CGGAATTCGAAGACTCAGTTTAACATTTATC-3′; Synpcc_term1R, 5′-CCCAAGCTTGGCTGGAAATCGCTG-3′; Synpcc_term2R, 5′-CCCAAGCTTGGCCGCACAGCTTC-3′.

### In Vitro Transcription Termination Assay

DNA templates for *in vitro* transcription were PCR amplified using gene blocks, as described above. DNA-RNAP complexes were assembled by incubating equal volumes of 2x DNA template (100 nM) and 2x reaction master mix for 5 min at 37°C. The master mix contained 400 nM cyRNAP core enzyme, 800 nM rpoD1, 100 µg/mL bovine serum albumin, and 2x transcription buffer (1x = 40 mM Tris-HCl, pH 8.0, 5 mM MgCl_2_, 5% glycerol, 0.1 mM EDTA, and 4 mM dithiothreitol [DTT]). RNAP and rpoD1 were added from a 10x stock solution containing 2 µM RNAP and 4 µM rpoD1 in enzyme dilution buffer (20 mM Tris-HCl, pH 8.0, 40 mM KCl, 1 mM DTT, and 50% glycerol). To the preformed DNA-RNAP complexes (2 µL aliquots), 1 µL of 4x protein mix containing NusA and/or NusG (final concentrations of each, 1.0 µM) was added, followed by incubation for 5 min at room temperature. Transcription was initiated by adding 1 µL of 4x rNTP mix (final 1x = 20 mM KCl, 250 µM each of ATP, GTP, and CTP, 50 µM UTP, and 0.5 µCi [α-^32^P] UTP in 1x transcription buffer). Reactions were incubated for 15 min at 37°C and terminated by adding an equal volume of 2x stop/loading buffer (40 mM Tris base, 20 mM Na_2_EDTA, 0.2% SDS, 0.05% bromophenol blue, and 0.05% xylene cyanol in formamide). RNA products were resolved on standard 5% polyacrylamide-urea sequencing gels, dried, and exposed using a phosphor screen. Signals were visualized using an Amersham Typhoon Phosphorimager, and band intensities were quantified using ImageJ to calculate the percent transcription termination (%T) as the ratio of terminated product to terminated and run-off product. The *in vitro* transcription termination experiment was performed twice, with each replicate derived from an independent reaction mixture. A representative gel is shown in **Supplementary Fig. 2**.

### TransTermHP Intrinsic Terminator Prediction for Defined and Diffuse Ends

To perform intrinsic terminator prediction, TransTermHP Version 2.07^23^ was applied to the *Syn* genome sequence without genome annotations and a confidence cutoff of 60 (-c 60). The prediction was performed separately on the forward and reverse strands of the genome sequence, and then the outputs from each run were combined for subsequent analysis.

Next, we determined the number of genes in each end category that had a predicted high-confidence intrinsic terminator in the downstream intergenic region. The intergenic region was defined as the sequence between the stop codon of the gene under consideration and the start codon of the next gene on the same strand. To stay within a biologically relevant range, terminator searches were limited to the first 1 kb if the intergenic region exceeded that length. If multiple intrinsic terminators were detected in the search region, we retained the terminator with the highest confidence score. After identifying genes with a predicted intrinsic terminator in the search region, we determined the number of genes in each end category with a high-confidence terminator by filtering for those with a confidence score > 80. Our cutoff is slightly more stringent than that used to define TransTermHP high-confidence terminators (confidence score ≥ 76)^23^.

### Mapping Transcript Tapering at Diffuse Ends

For diffuse ends, our pipeline identifies the position where read density drops at least ∼70% (3.5-fold) relative to the region upstream of the stop codon and records this as the diffuse end position. To estimate the tapering length of mapped reads as they ramp down at diffuse transcript termini, we measured the distance from the stop codon to this endpoint (“post-stop taper length”) in **Fig. 4a**. The read counts from WT diffuse ends were also used to generate the diffuse end traces in **Fig. 4b**. For each gene, the read density was calculated in 100 bp windows spaced 15 bp apart, starting 513 bp upstream of the center of the 75 bp window where the density drop was identified and extending 262 bp downstream. The read densities for each window were then normalized to the read density in the gene body (calculated across a window extending 30-105 bp upstream of the stop codon). Diffuse ends with multiple 3′ peaks (diffuse peaks, n=75) were excluded from this analysis. A randomly selected subset of 10 diffuse ends is shown in the main text; all analyzed diffuse ends can be seen in **Supplementary Fig. 3**.

### Sequence Bias Analysis at TU Ends and TU-Internal Gene Ends

To analyze the sequence composition around *Syn* defined and diffuse ends, the fractions of A, T, C, and G nucleotides were determined in windows extending 51 or 201 bp upstream and downstream of (1) the 3′ peak for all the identified defined ends (n=388), (2) the downstream end of the window corresponding to the density drop for diffuse ends (excluding diffuse peaks) (n=274), and (3) the downstream end of the final window reached by the end calling pipeline for all *Syn* genes in TUs (i.e., TU-internal genes) (n=544). Statistical comparisons were performed using the unpaired two-sided t-test (scipy.stats.ttest_ind) with unequal variance assumed (equal_var=False).

### Synechocystis RNase J Depletion Analysis

To compare the TU end positions in RNA-seq data from WT and RNase J-depleted *Synechocystis* sp. PCC 6803^82^, the end calling pipeline (see above section) was modified to eliminate use of 5′ and 3′ peak information. Briefly, the modified pipeline was used to scan for a drop in read density exceeding 4-fold between the gene body (30-105 bp upstream of the stop codon) and a sliding 75-bp window that could extend as far as the stop codon of the following gene on the same strand. The end position of the window where the read density first dropped below 4-fold was recorded. Classification was performed using the gene annotations corresponding to NC_000911.1, excluding genes shorter than 155 bp or with fewer than 5 reads/bp across the gene body (excluding 45 bp on either end).

To determine if TU ends tended to shift downstream in the RNase J depletion strain, we calculated the positional offset between the TU end identified in WT and RNase J-depleted *Synechocystis* (Position_depletion_ – Position_WT_). Positive offsets indicate that the drop in density was observed farther downstream in the depletion dataset. The data shown in **Fig. 5** are limited to genes that were identified as terminated in the WT data and exceeded 100 reads/bp in the gene body.

### BLAST Searches for Termination Factor Homologs in Cyanobacteria

To test for cyanobacterial homologs of known archaeal, eukaryotic, and organellar termination factors, the following BLAST searches were performed: (1) a protein-protein BLAST (blastp) of *A. thaliana* RHON1 (GenBank: OAP15819.1) against the *S. elongatus* PCC 7942 (taxid:1140) non-redundant protein sequences (nr) database, and an expanded blastp search against the entire cyanobacterial phylum (taxid:1117) nr database; (2) blastp of MTERF1 from *Homo sapiens* (UniProtKB/Swiss-Prot: Q99551.1) and *Mus musculus* (isoform X1, NCBI RefSeq: XP_017176489.1) against the cyanobacterial (taxid:1117) and *S. elongatus* PCC 7942 (taxid:1140) nr databases, and (3) blastp and Domain Enhanced Lookup Time Accelerated BLAST (DELTA-BLAST) of FttA from a representative hyperthermophilic archaeon *Pyrococcus abyssi* GE5^97^ (UniProtKB/Swiss-Prot: Q9V0P0.2) against the *S. elongatus* PCC 7942 (taxid:1140) nr database.

### Displacement and Readthrough Fold-Change Calculations for mfd Knockout

To assess the impact of *mfd* knockout on TU end positions, we calculated the positional offset of TU ends in each end class between WT and the Δ*mfd* strain (Position_Δ*mfd*_ – Position_WT_). Biological reproducibility between Δ*mfd* replicates was similarly assessed by computing the offset between replicates (Position_Δ*mfd* rep 2_ – Position_Δ*mfd* rep 1_). TU end positions were defined as either the 3′ peak position (for defined ends) or the end of the downstream window where the density drop threshold is met (for diffuse ends), as determined by the end calling pipeline.

To quantify the effect of *mfd* knockout on transcriptional readthrough (RT) at defined ends, we used the following log2-based ratios:

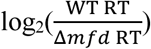

to compare readthrough levels between WT and Δ*mfd* replicates, where negative values on the x-axis indicate increased readthrough in the Δ*mfd* strain, and

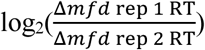

to evaluate biological reproducibility between knockout replicates. RT values were calculated using the end calling pipeline.

### Data Availability

All sequencing data (raw FASTQ and shadow-removed WIG files) have been deposited in the Gene Expression Omnibus under accession number GSE309256.

## Acknowledgments

The authors would like to thank the Li lab for useful discussion and interpretation, especially James Xue for support designing the RNA extraction protocol and performing shadow-removal and Mirae Parker for help establishing the peak-calling method, and the Chisholm lab for guidance on cyanobacterial cell culturing. G.W.L. is an investigator at the Howard Hughes Medical Institute. This research was supported by NIH R35GM124732 (to G.W.L.), R35GM156623 (to K.S.M.), R35GM153190 (to P.B.), UKRI BBSRC Mission Award BB/Y007638/1 (to Y.Y.), the Pew Innovation Fund (to G.W.L. and K.S.M.), MathWorks graduate fellowship (to J.A.C. and K.J.D.), and NIH T32GM136540 (to J.A.C. and K.J.D.).

## Contributions

J.A.C., K.J.D., and G.W.L. conceptualized the research. J.A.C. generated the Mfd knockout strain and performed all *in vivo* experimental work. Y.Y. provided cyNusA and cyNusG expression vectors. R.K.V., P.B., and K.S.M. designed and performed the *in vitro* experiment. J.A.C. developed experimental methodology, and K.J.D. wrote the end calling pipeline. J.A.C. and K.J.D. performed data analysis and prepared all data visualizations. G.W.L., K.S.M., P.B., and Y.Y. acquired funding, and G.W.L. provided resources and supervision. J.A.C wrote the original manuscript draft, and J.A.C., K.J.D., P.B., and G.W.L. edited the manuscript. All authors reviewed the manuscript.

## Supplementary Figures

**Supplementary Figure 1.**
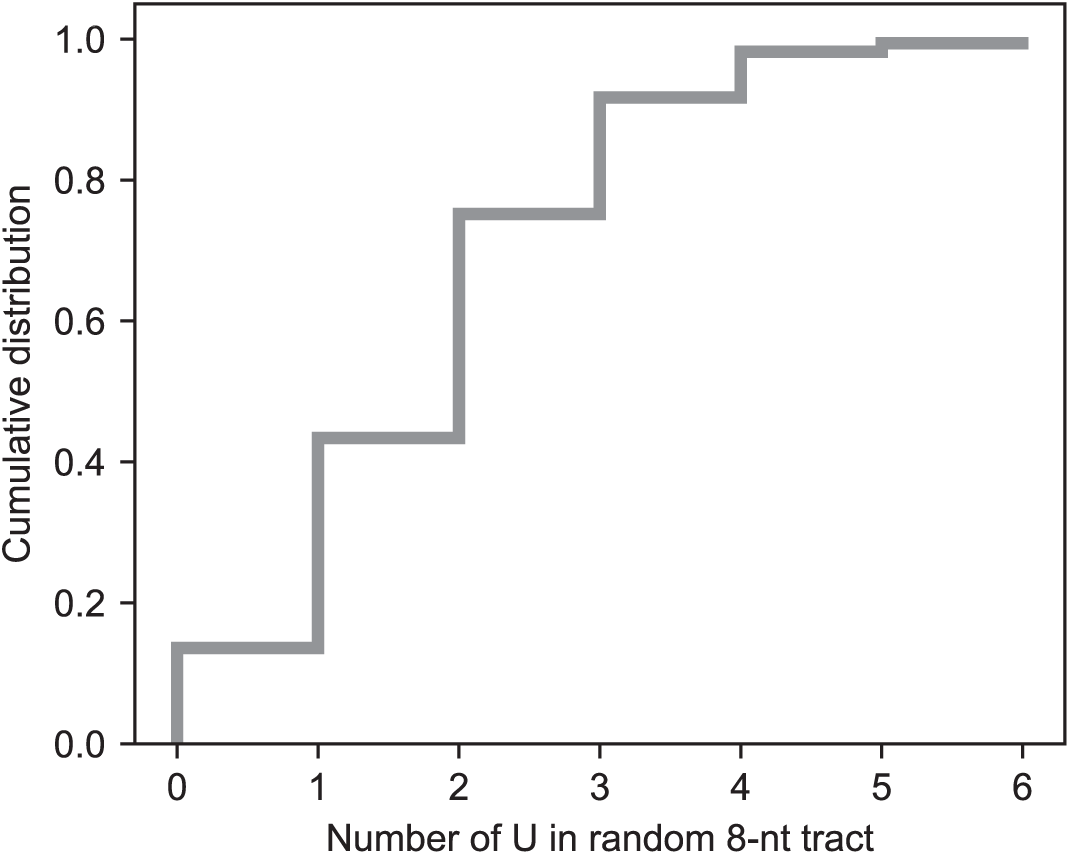
Cumulative distribution of uridine counts in 1,000 random 8-nucleotide (nt) sequence tracts in *Synechococcus elongatus* PCC 7942. Sequences were sampled evenly from the forward and reverse strands. A threshold of ≥ 4 uridines (8% chance of occurring at random) was chosen to define U-tracts. There is a 2% chance of ≥ 5 uridines occurring at random in an 8-nt tract and a 25% chance of ≥ 3 uridines.

**Supplementary Figure 2.**
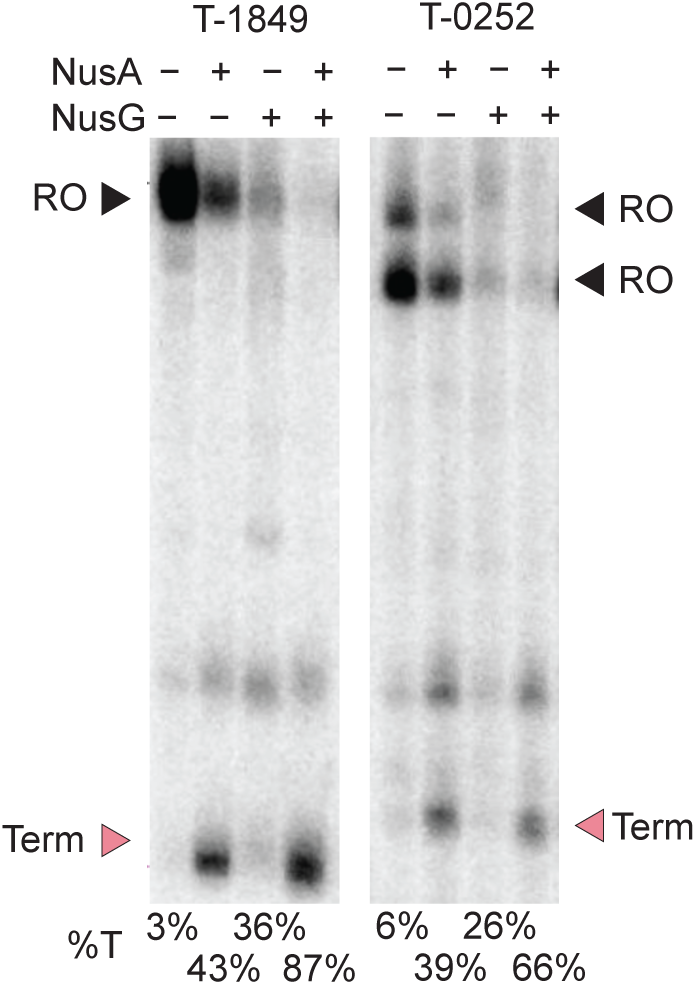
Multi-round *in vitro* transcription termination assays for the intrinsic terminators downstream of the genes *Synpcc7942_1849* (left) and *Synpcc7942_0252* (right). Experiments were performed in the presence (+) or absence (-) of NusA and/or NusG as indicated (**Materials and Methods**). Positions of terminated (Term, pink arrow) and run-off (RO, black arrow) product are marked. Termination efficiencies (%T) for each terminator under the specified conditions are indicated below the lanes (**Materials and Methods**).

**Supplementary Figure 3.**
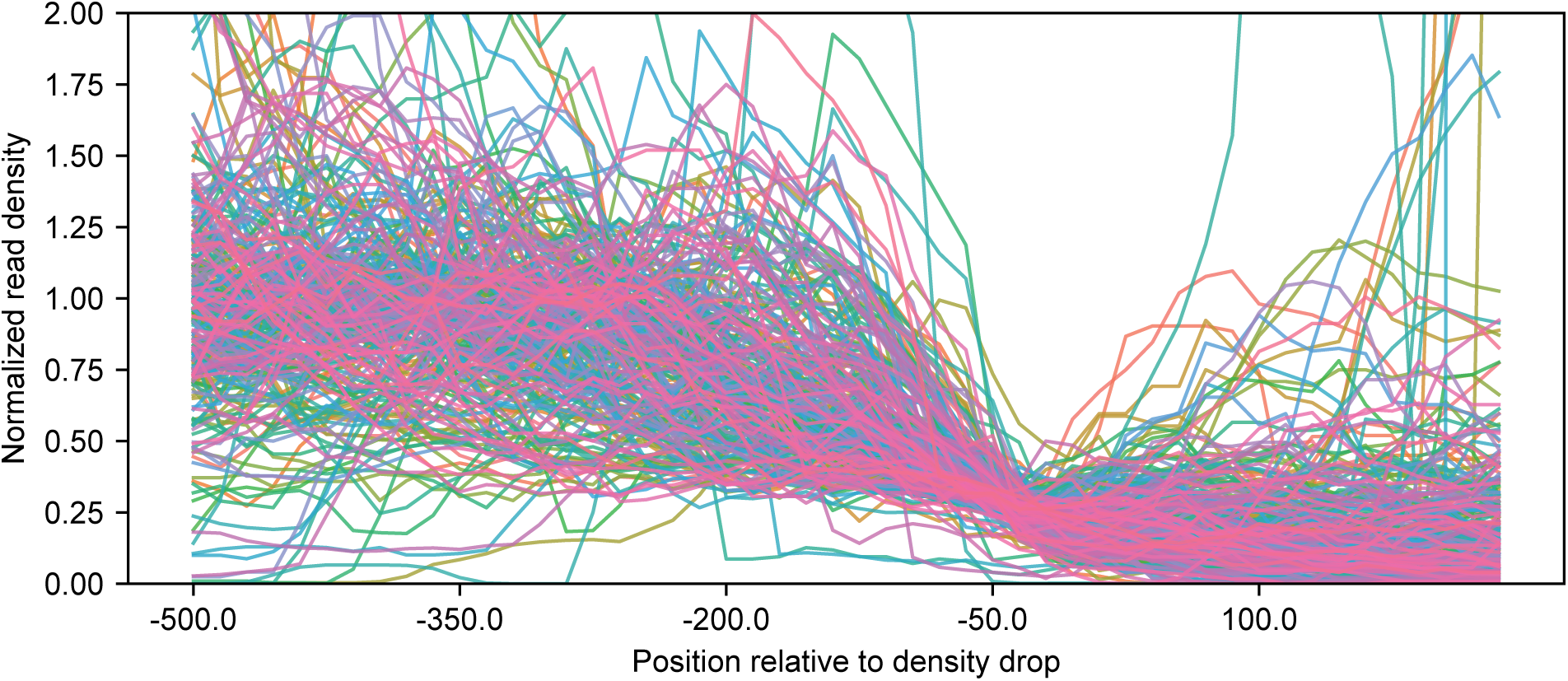
Global tapering behavior at *Synechococcus elongatus* PCC 7942 diffuse TU ends. Normalized read densities were calculated in a fixed window around diffuse end positions as described in **Materials and Methods**. All diffuse ends are shown except those that contain multiple 3′ peaks (diffuse peaks) (n=274).

**Supplementary Figure 4.**
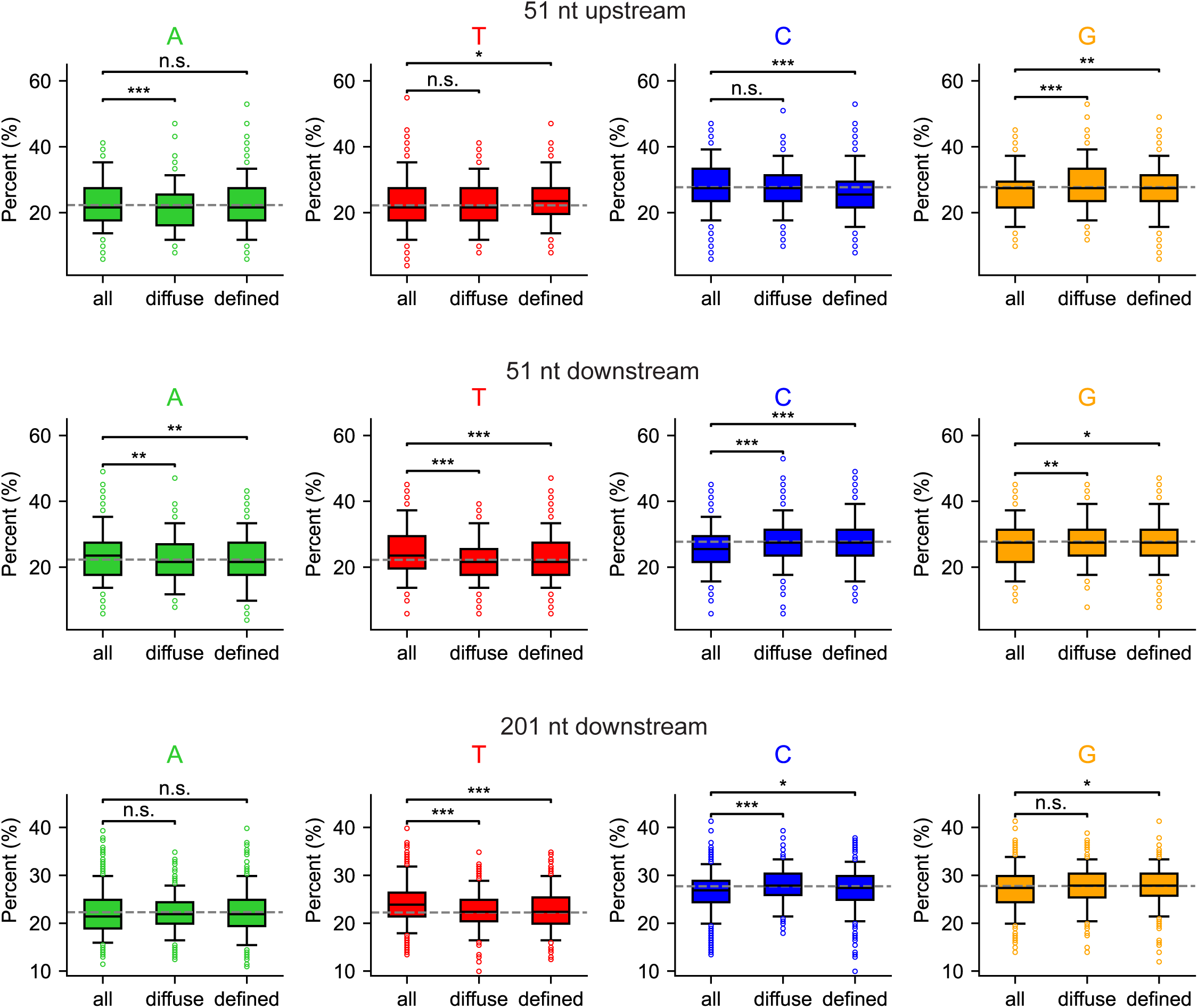
Nucleotide (nt) composition analysis in upstream 51-nt window and downstream 51-nt and 201-nt windows flanking diffuse TU ends in *Synechococcus elongatus* PCC 7942. Nucleotide content in the specified windows was computed for defined and diffuse TU ends (“defined”, “diffuse”) and the 3′ ends of all internal operon genes (i.e., within TUs but not at TU 3′ end), used to control for general gene-end effects (“all”). Box plots show A/T/C/G content distributions; whiskers represent the 5^th^ and 95^th^ percentiles. Gray dotted lines indicate background genomic nucleotide frequencies. Statistical significance was assessed using unpaired two-sided Welch’s t-tests (n.s. = not significant; * = p < 0.05; ** = p < 0.01; *** = p < 0.001). Biases observed in the 51-nt upstream window mirror those in the 201-nt upstream window but are less pronounced. No clear biases were detected downstream of diffuse TU ends.

**Supplementary Figure 5.**
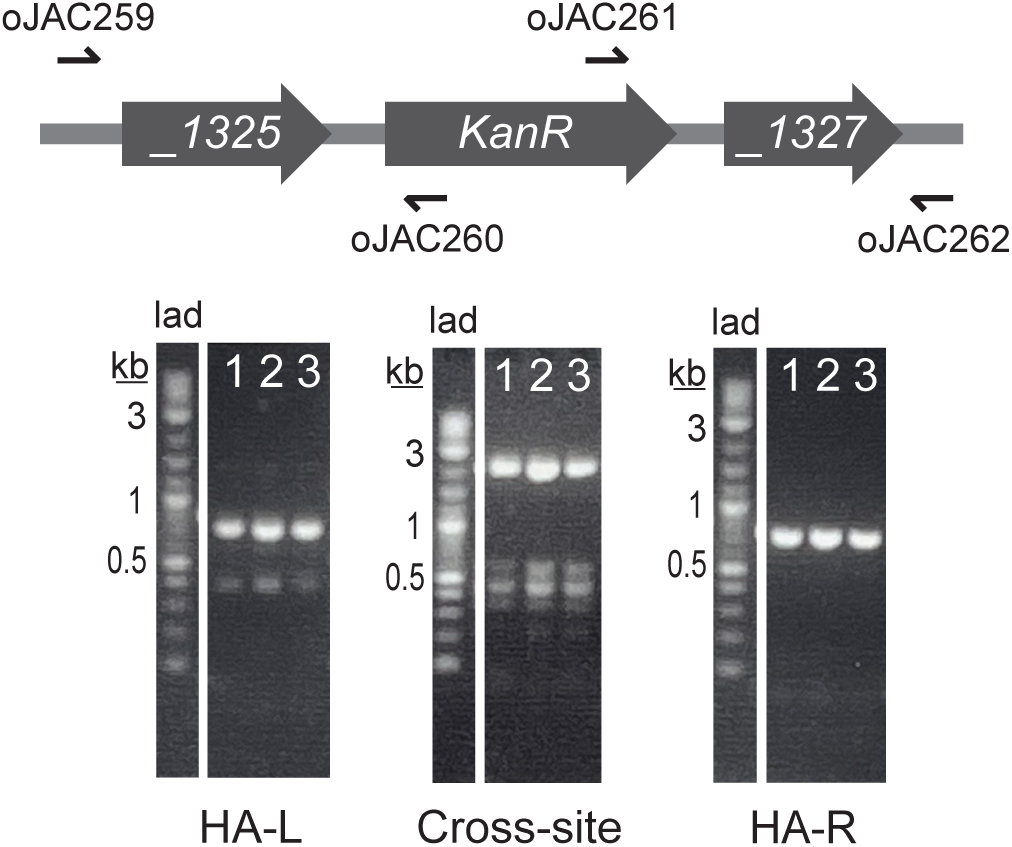
Chromosomal PCR confirms Δ*mfd* construction. PCR product sizes are as expected (see **Materials and Methods**) for the left junction, right junction, and cross-site reaction on the three tested colonies, confirming successful knockout strain construction. Colony 3 was selected as the strain representative for use in Rend-seq. HA-L = left homology arm junction; HA-R = right homology arm junction; *KanR* = kanamycin resistance gene (*aphI*); *_1325* = *Synpcc7942_1325*; *_1327* = *Synpcc7942_1327*; lad = Quick-Load® 1 kb Plus DNA Ladder (NEB, Catalog No. N0469S).

**Supplementary Figure 6.**
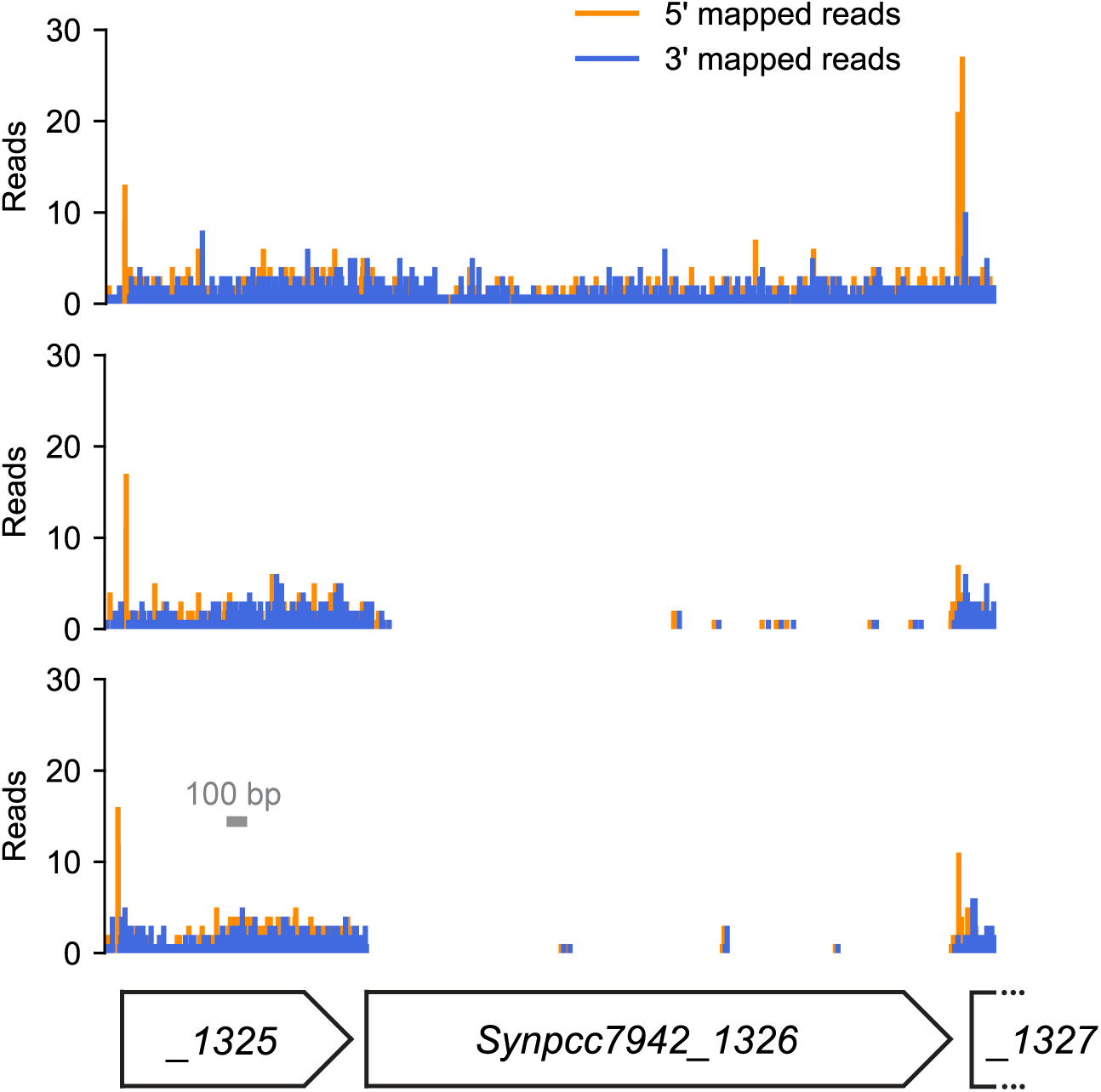
Mfd knockout confirmed at the RNA level in *Synechococcus elongatus* PCC 7942 (*Syn*). Rend-seq traces for *Synpcc7942_1326* (*mfd*) and the surrounding genes demonstrate successful transcript-level knockout in Δ*mfd* replicates (reps). Top trace corresponds to wild-type *Syn*, middle trace is Δ*mfd* rep 1, and bottom trace is Δ*mfd* rep 2. Gray bar shows x-axis scale. *_1325* = *Synpcc7942_1325*; *_1327* = *Synpcc7942_1327*.

**Supplementary Figure 7.**
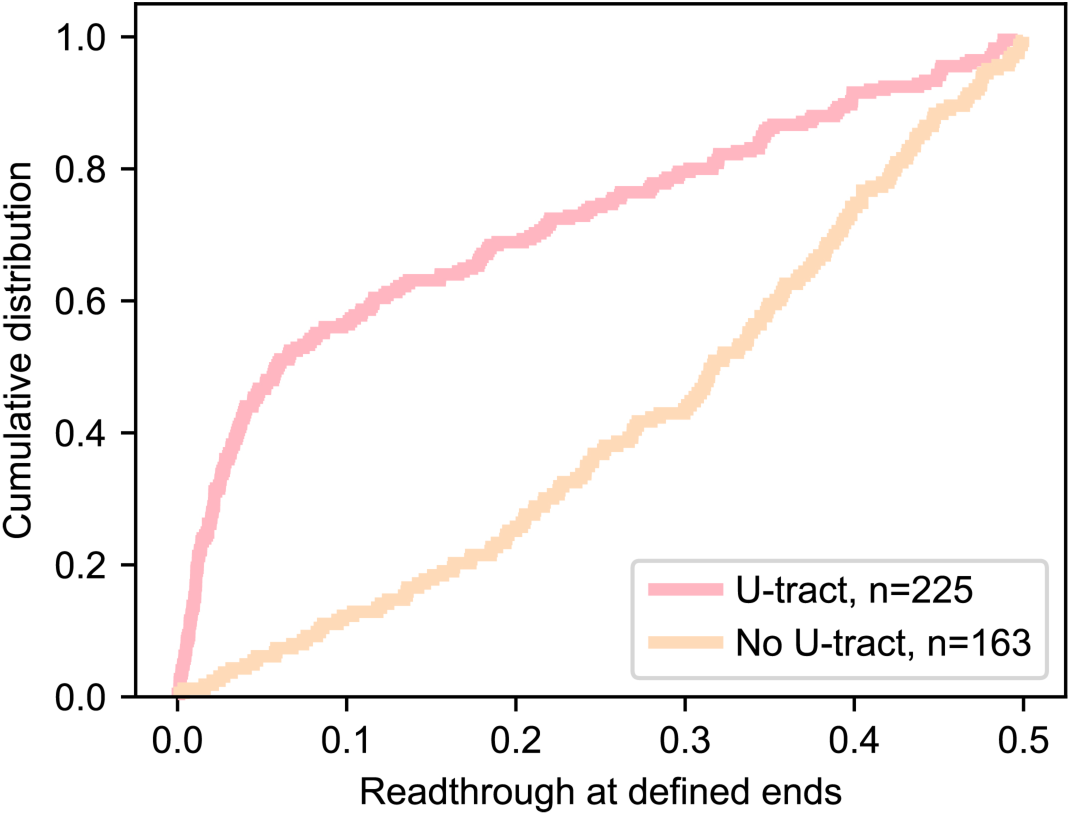
Transcriptional readthrough at defined ends with and without U-tracts. Cumulative distributions of readthrough values (calculated as described in **Materials and Methods**) show elevated baseline readthrough at defined ends lacking U-tracts compared to those with U-tracts.

